# People with nicotine use-disorder exhibit more prefrontal activity during preparatory control but reduced anterior cingulate activity during reactive control

**DOI:** 10.1101/2021.12.08.471840

**Authors:** Shivam Kalhan, Li Peng Evelyn Chen, Marta I. Garrido, Robert Hester

## Abstract

Reduced inhibitory control and a hypersensitivity to reward are key deficits in drug-dependents, however, they tend to be studied in isolation. Here we seek to understand the neural processes underlying control over reward and how this is different in people with a nicotine use disorder (pNUD). A novel variant of the monetary incentive delay task was performed by pNUD (n = 20) and non-smokers (n = 20), where we added a stop-signal component such that participants had to inhibit prepotent responses to earn a larger monetary reward. Brain activity was recorded using functional magnetic resonance imaging (fMRI). We estimated stop signal reaction times (SSRT), an indicator of impulsivity, and correlated these with brain activity. Inhibitory accuracy scores did not differ between the control group and pNUD. However, pNUD had slower SSRTs, suggesting that they may find it harder to inhibit responses. Brain data revealed that pNUD had greater preparatory control activity in the middle frontal gyrus and inferior frontal gyrus prior to successful inhibitions over reward. In contrast, non-smokers had greater reactive control associated with more activity in the anterior cingulate cortex during these successful inhibitions. SSRT-brain activity correlations revealed that pNUD engaged more control related prefrontal brain regions when SSRTs are slower. Overall, whilst the inhibition accuracy scores were similar between groups, differential neural processes and strategies were used to successfully inhibit a prepotent response. The findings suggest that increasing preparatory control in pNUD may be one possible treatment target in order to increase inhibitory control over reward.

## INTRODUCTION

Addiction is a complex disorder which includes multiple decision-making symptoms^1^. Key deficits include hypersensitivity towards drug-related rewards^2^, including towards monetary rewards^3^, and reduced inhibitory control^4^. These deficits may contribute to the high persistence in drug-seeking behaviors and impulsive decision-making found in people with a substance use disorder (pSUD). Impulsivity is defined here as overvaluing a smaller immediate drug-reward compared to the larger later reward of better health from abstinence^5^. A better understanding of these processes may offer insight into ways of decreasing drug-seeking behaviors and avoiding relapse by exercising control over immediate rewards. These differential processes of inhibiting a response and the hypersensitivity to rewards have been relatively well studied; but are typically studied independently.

The stop-signal task is commonly used to study processes involved in response inhibition^6^. In this task, there is a prepotent tendency to respond to a target stimulus due to the frequent “go-trials”. However, less frequently, there is a stop-signal presented with a short latency to the target stimulus, where individuals now need to withhold their response. These are termed “stop-trials”. The dorsolateral prefrontal cortex (dlPFC), including the inferior frontal gyrus (IFG) are associated with successfully inhibiting a response during stop-trials^7^, both, prior to inhibition (preparatory control^8,9^) and during inhibition (reactive control^10^). Those with a SUD, including to nicotine, have demonstrated reduced inhibitory control in this stop-signal task^11–13^ and this has been associated with hypoactivity in regions within the PFC^14^.

The mechanisms underlying reward-processing have been studied through various paradigms, including the monetary incentive delay (MID) task^15^. The MID task cues participants on possible upcoming rewards and then asks them to respond to a target stimulus to win or lose rewards – depending on the variant of the task used. The striatum has typically been associated with reward processing^16^ and in this MID task, increased striatal activation has been associated with reward anticipation^17^. Administering the MID task to participants with a SUD has identified aberrant striatal functioning (with increases and decreases relative to controls), particularly when they are asked to anticipate rewards. This aberrant striatal functioning is thought to contribute to the heightened impulsivity and impaired reward processing^18^ and although the opponent-process theory^19^ suggests that this striatal aberration is a possible predictor of developing addictive-like behaviors, evidence supporting this relationship is mixed^20,21^. Lastly, PFC hypoactivation during reward processing is also found in pSUD, implying that there may be reduced inhibitory control over choices with immediate rewards^22^.

Taken together, the PFC appears to increase cognitive control and the striatum plays a role in processing and anticipating rewards. Both of these brain regions exhibit aberrant functioning in addiction and possibly contribute to the reduced inhibitory control over immediate reward which may drive impulsive decision-making. However, the neural mechanisms underlying how the two processes interact remains less clear. Namely, how does the brain exercise inhibitory control over reward in pSUD? Is this through modulation of PFC activity, striatal activity, or both? We hypothesized that reduced PFC activity may be associated with failed inhibitions, consistent with its role in cognitive control. We further hypothesized that increased striatal activity may also be associated with failed inhibitions, by possibly anticipating and over-valuing the smaller sooner rewards.

To investigate this question, we used a novel variant of the MID task – termed monetary incentive control task (MICT). Here, we added the stop-signal component to the MID task. We also used real monetary reward components of two modalities; smaller sooner and larger later. To win the smaller reward, participants had to quickly respond to a target stimulus. To win the larger reward, participants had to successfully inhibit their response on most of the stop-trials. If participants responded quickly on stop-trials – they still won the small reward. Previous studies^23,24^ have also combined the stop-signal task with a reward manipulation, however, a critical distinction is that in our task there is also a reward given for failing to inhibit, whereas reward was only given for successfully inhibiting in the previous tasks. Failed inhibition was rewarded to better simulate real-world abstinence where failing to inhibit comes with a small immediate reward. Importantly, participants were not updated on their progress towards winning the larger reward, since again, the aim was to simulate real world abstinence and attempting abstinence does not come with the immediate certainty of the receipt of a larger reward (better health in the future). However, a relapse comes with the more certain small reward (drug).

Previous accounts have found that monetary incentives can 1) increase reactive control in the stop-signal task by increasing engagement of control-related PFC activity^25^, 2) reduce conflict during a conflict-response task, associated with increased functional connectivity between intraparietal sulcus and the striatum, engaging more top-down control^26^ and, 3) increase performance in the Stroop-task in both healthy controls (by enhancing dorsolateral PFC activity) and in people with a cocaine use-disorder (by enhancing occipital lobe activity and the functional connectivity between the dorsolateral PFC and striatum)^27^. In contrast, our task provides a smaller sooner incentive for failed inhibitions, and accordingly, we hypothesized that this may make inhibitions more difficult by engaging striatal anticipatory activity, particularly in pSUD, and successful inhibitions may require increased PFC engagement, especially to exercise control over the smaller sooner incentive for failed inhibitions. There was also a manipulation where participants were cued on the probability that the upcoming trial would be a ‘stop’ trial. This manipulation was used to investigate the potential interaction between reward anticipation and ‘stop’ difficulty, with a higher probability condition hypothesized to engage control related PFC regions and a higher stop accuracy score.

We recorded brain activity using functional magnetic resonance imaging (fMRI) while participants performed this task. Both groups of people with a nicotine use disorder (pNUD) and non-smokers healthy controls underwent this task. Nicotine withdrawal can produce neurotoxic effects in the mesolimbic reward systems comparable to other drugs of abuse including amphetamine, cocaine and opiates^28^. We also estimated stop signal reaction times (SSRTs), which is a measure of how effortful it is to inhibit a response, as well as an index of impulsivity^29^. Overall, we investigated how the brain may exercise control over smaller sooner rewards to attain a larger later reward in pNUD and non-smokers.

## MATERIALS AND METHODS

### Participants

Participants were recruited through advertisements at the University of Melbourne, and through a community website. All participants provided written informed consent which was approved by Human Ethics Committee of the University of Melbourne and the Royal Children’s Hospital. The pNUD group consisted of twenty individuals (10 males, 10 females; mean age = 24.3 years, standard deviation = 4.7, range = 18 – 34). The brain structural images for one pNUD group participant was not retrievable, hence brain imaging data for this participant was not analyzed. The control group consisted of twenty non-smokers (10 males, 10 females; mean age = 23.7 years, standard deviation = 4.3, range = 18 – 32). One participant in the control group had an anatomical anomaly, however, this was very minor, and the participant was included in the analysis. All participants were right handed – determined by the Edinburgh Handedness Inventory ^30^. Participants in the control group had smoked less than 6 cigarettes in their lifetime. Participants in the pNUD group smoked at least fifteen cigarettes daily and the average Fagerstrom Test for Nicotine Dependence (FTND) score was 3.95, indicating close to moderate dependence^31^. The group average for years of cigarettes smoked was 7.6 years.

Prior to the experiment, the pNUD group had been abstinent for at least three hours. This was confirmed by both a self-report and the carbon monoxide breath measure. Exclusion criteria for both groups consisted of a history of neurological or psychiatric disorder, current use of psychotropic medication (other than nicotine for the pNUD group). Scanning data was collected between 9am and 5pm across participants, and they were advised not binge prior to abstaining for the 3-hours. Participants also arrived at least 1-hour prior to scanning for task preparation/practice, which also prevented them from smoking. The 3-hour abstinence window was chosen due to nicotine’s half-life of approximately 2-hours^32^, and data suggesting that 3-hours abstinence did not produce withdrawal effects on cognition^13^.

### Monetary Incentive Control Task (MICT) Behavioral Paradigm

The MICT paradigm is a modified version of the monetary incentive delay (MID) task, but with the addition of a stop-signal component (see Figures 1 and 2). Each trial began with a cue symbol, presented for 2 s. The cue was used to inform participants on the probability the upcoming trial was a stop trial and whether it is a reward or a neutral trial. Following this cue epoch, there was a variable delay presented for 2 – 4 s, with 1 s jitter. This delay period was termed the anticipation epoch and comprised of a blank screen. After this, the target was presented. This was either a “X” or an “O”, presented for 400ms, with a blank screen for 600ms following this. Participants had to press the correct button (left or right) associated with the target letter within 400ms to get the small 20¢ reward, if this was a reward trial. For stop trials, a square border around the target letter would appear after 150ms latency. If this stop trial was a reward trial and participants withheld their response at 60% or more of these rewarding stop trials, they would get the large $20 reward at the end of the task. If they responded within 400ms to the rewarding stop trial – they would still get the small 20¢ reward. For neutral trials – no monetary reward could be received, irrespective of performance. Overall, there were four conditions based on the following trial types - 1) reward trial with 20% probability (R20), 2) neutral trial with 20% probability (N20), 3) reward trial with 40% probability (R40) and 4) neutral trial with 40% probability (N40). These could all be either “go” or “stop” trial types. Following the trial – participants were presented with feedback for 1.5 s. The feedback indicated their performance and if they had won the small 20¢ reward (see Figures 1 and 2). Following each run of trials, participants were given feedback on how much money they had earned from go-trials. However, importantly, feedback on stop trial accuracy and any associated monetary gain on the larger $20 reward was not provided until the end of the task.

**Figure 1.**
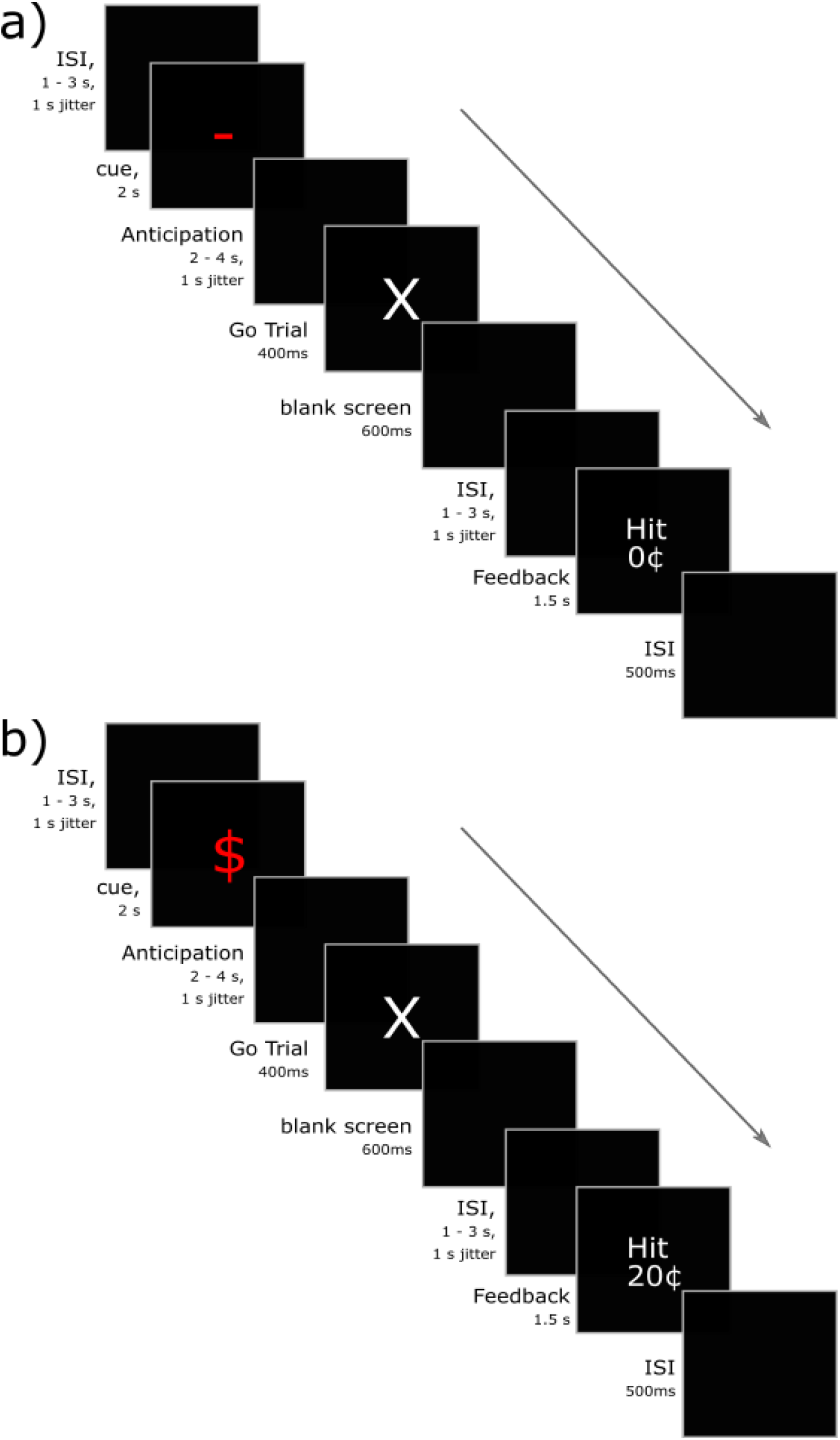
Example sequences of go-trials. If the cue symbol was a dash (“-“), the upcoming trial was a neutral trial and no money could be won, irrespective of performance. If it was a dollar symbol (“$”), the upcoming trial would be a reward trial where monetary rewards could be won, depending on performance. If the color of the symbol was white – there was a 20% probability that the upcoming trial would be a stop trial. If this color was red – it would be 40% probability. **a)** N40 go-trial which is accurately performed (correct button-press under 400ms). Feedback is “hit” but because this is a neutral trial, no monetary reward is won. **b)** R40 go-trial which is accurately performed (correct button-press under 400ms). Feedback is “hit” and because this is a reward trial, a 20¢ reward is won.

**Figure 2.**
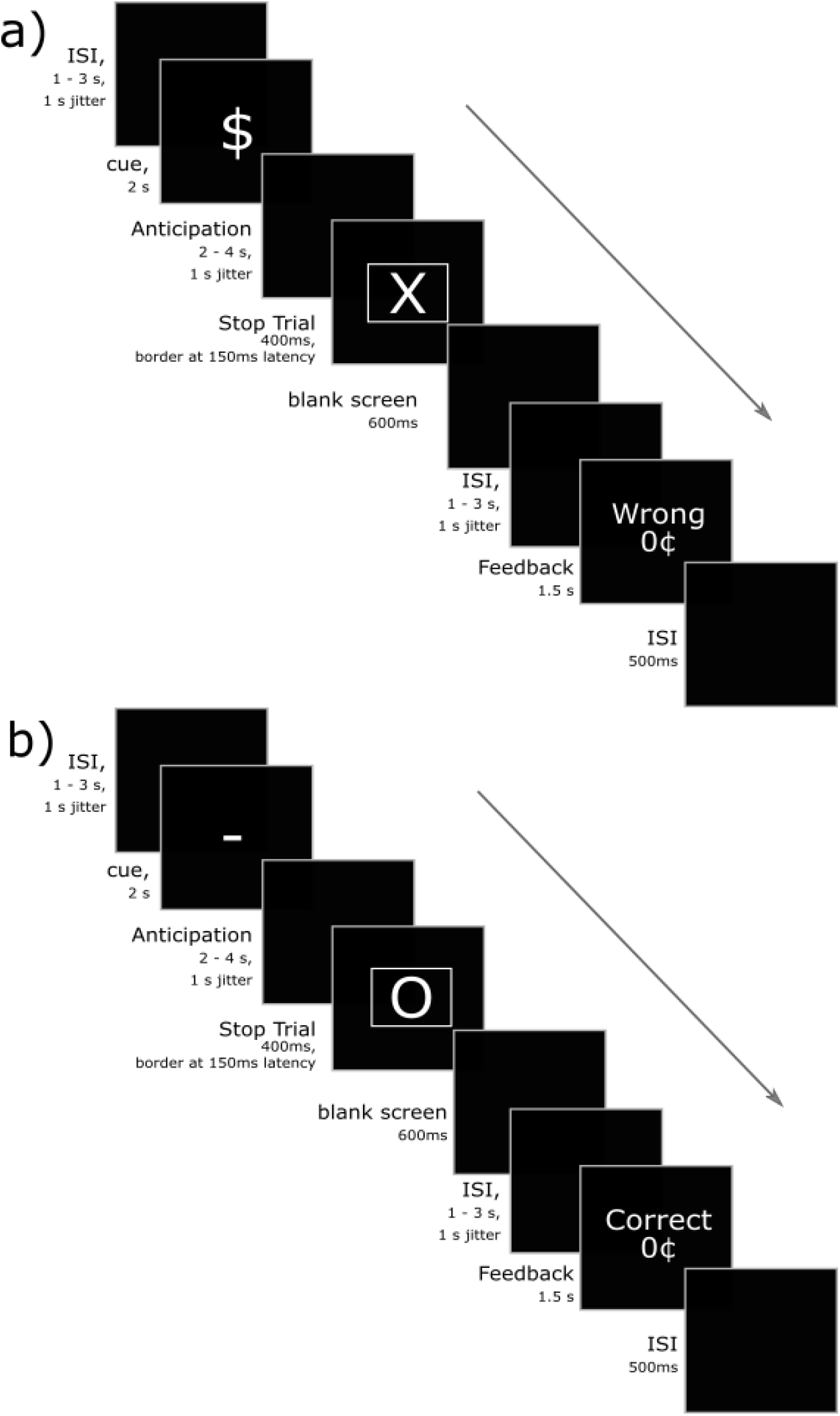
Example sequences of stop-trials. If the cue symbol was a dash (“-“), the upcoming trial was a neutral trial and no money could be won, irrespective of performance. If it was a dollar symbol (“$”), the upcoming trial would be a reward trial where monetary rewards could be won, depending on performance. If the color of the symbol was white – there was a 20% probability that the upcoming trial would be a stop trial. If this color was red – it would be 40% probability. **a)** R20 stop-trial with the stop signal appearing after 150ms latency. In this example, the participant failed to inhibit a response and did not respond within the 400ms. Feedback “wrong” is given for responding (failed inhibition) and no reward is given as the response wasn’t under 400ms. **b)** N20 stop-trial with the stop signal appearing after 150ms latency. In this example, the participant accurately inhibited their response (successful inhibition).

There was a total of 216 trials, with 54 of these being stop trials (25%). This different break-up of ‘stop’ and ‘go’ trials (25% stop-trials and 75% go-trials) is commonly used to create the prepotent tendency to respond in the stop-signal paradigm^6^. The task was split into six runs, with 36 trials per run. Trials were presented in a pseudo-random, intermixed design, within each run. The R20 condition had a total of 78 go-trials and 18 stop-trials, R40 had 30 go-trials and 18 stop-trials, N20 39 go-trials and 9 stop-trials, N40 had 15 go-trials and 9 stop-trials. The task lasted approximately 45 minutes including rest breaks between runs. Please see supplementary material methods for 1) the apparatus details, 2) experimental procedure, 3) MRI sequences used and 4) the methods of SSRT estimation.

### Behavioral Analysis

For performance indices we used the stop-accuracy percentage (calculated as the number of stop-trials where participants inhibited their responses divided by the total number of stop-trials) and go-accuracy percentages (calculated as the number of responses made on go-trials in under 400ms divided by the total number of go-trials). Data above and/or below three standard deviations from the mean were removed as outliers. To test for significance, we used repeated-measures analysis of variance (ANOVA). There were three factors and each factor had two levels (2 x 2 x 2 ANOVA design). These included; 1) factor of probability, with levels of low (20%) and high (40%) probabilities, 2) Factor of reward, with levels of rewarding trial and neutral trial, and 3) factor of group, with levels of pNUD group and control group. To test for simple effects, independent t-tests were conducted where ANOVA yielded significant results, corrected for multiple comparisons using Šidák correction. The partial eta-squared (η_p_^2^) was calculated as a measure of effect size with 0.01 being small, 0.06 being medium and 0.14 being large.

### fMRI Analysis

Images were pre-processed using SPM default functions with steps in the order of realignment, co-registration, segmentation and normalized into MNI-space. The images were smoothed using the 6mm FWHM Gaussian kernels. Runs with movement of greater than 4mm (one voxel size) in either of the x, y, z planes were removed from further analysis. All general linear models (GLMs) consisted of six movement regressors. A high-pass filter of 128s was used.

Only the stop-trials were analyzed, across all conditions and for both failed or successful inhibitions. This gave eight regressors of interest in our first-level GLM (four conditions for failed and successful inhibitions). These were all analyzed for four different epochs; 1) cue epoch, 2) anticipation epoch, 3) trial epoch (from onset of stop signal) and 4) feedback epoch. We modelled cue and anticipation epochs separately to investigate any possible brain activity differences during encoding of the cues, which are only presented during the cue epoch and not during the anticipation epoch. All task epochs apart from the epoch of interest was modelled as a regressor of no interest. For example, when analyzing for effects at the stop trials for the cue epoch – all other epochs at these stop trials were modelled as regressors of no interest, in addition to also all epochs in go-trials. This was done to reduce noise at our epoch and trial of interest, while reducing possible confounds from other trials and epochs. Lastly, for the feedback epoch, failed inhibitions where a 20c reward was won, was excluded. This was to remove any potential confound of the small 20c reward and to directly compare feedback of 1) “correct 0c” for successful inhibitions and “miss 0c” for failed inhibitions.

Second-level models were performed using a full-factorial, 2 x 2 x 2 ANOVA design. This included three factors, with two levels each; 1) group (control and pNUD), 2) inhibition accuracy (failed and successful) and 3) reward (neutral trial or reward trial). Main effects and interactions were tested and, t-contrasts to examine condition-specific results. For model-based results, SSRTs for each participant, across each condition were aligned and used as covariates of interests at the second-level. This GLM consisted of cells for each reward and probability condition, for each of the two groups. There was no inhibition accuracy factor in this GLM as SSRT is not estimated for failed inhibitions (where a response is made). All data presented has threshold of p < 0.05 family-wise error corrected (FWE) at the cluster-level, unless specified otherwise. Background brain image used for figures is from SPM canonical, which is an average T1 from 305 individuals, in MNI-space.

## RESULTS

### Behavior

#### pNUD have similar stop accuracy scores to controls

As in Figure 3a, there were no significant differences in stop accuracy scores between groups (main effect of group: F(1,38) = 3.2, p = 0.08, η_p_^2^ = 0.078), and no main effect of reward (F(1,38) = 2.3, p = 0.13, η_p_^2^ = 0.058). There was a main effect of probability (F(1,38) = 65.95, p < 0.0001, η_p_^2^ = 0.63), with performance better in the 40% probability condition compared to the 20%. There was also a significant group x probability interaction (F(1,38) = 4.70 p = 0.036, η_p_^2^ = 0.11), which appeared to be driven by the differences between pNUD and control groups in the N20 condition (t-test, p = 0.007). Twelve participants from the pNUD group won the larger later $20 reward, and thirteen participants from the control group. This was won by successfully inhibiting a response on 60% or more of the reward stop-trials. Overall, the stop-accuracy scores suggest that both groups perform better when cued that the upcoming trial has a higher probability of being a stop-trial, and this is irrespective of whether this is a reward or a neutral trial. See Figure S1 (*supplementary materials*) for accuracy scores and reaction times for go-trials.

**Figure 3.**
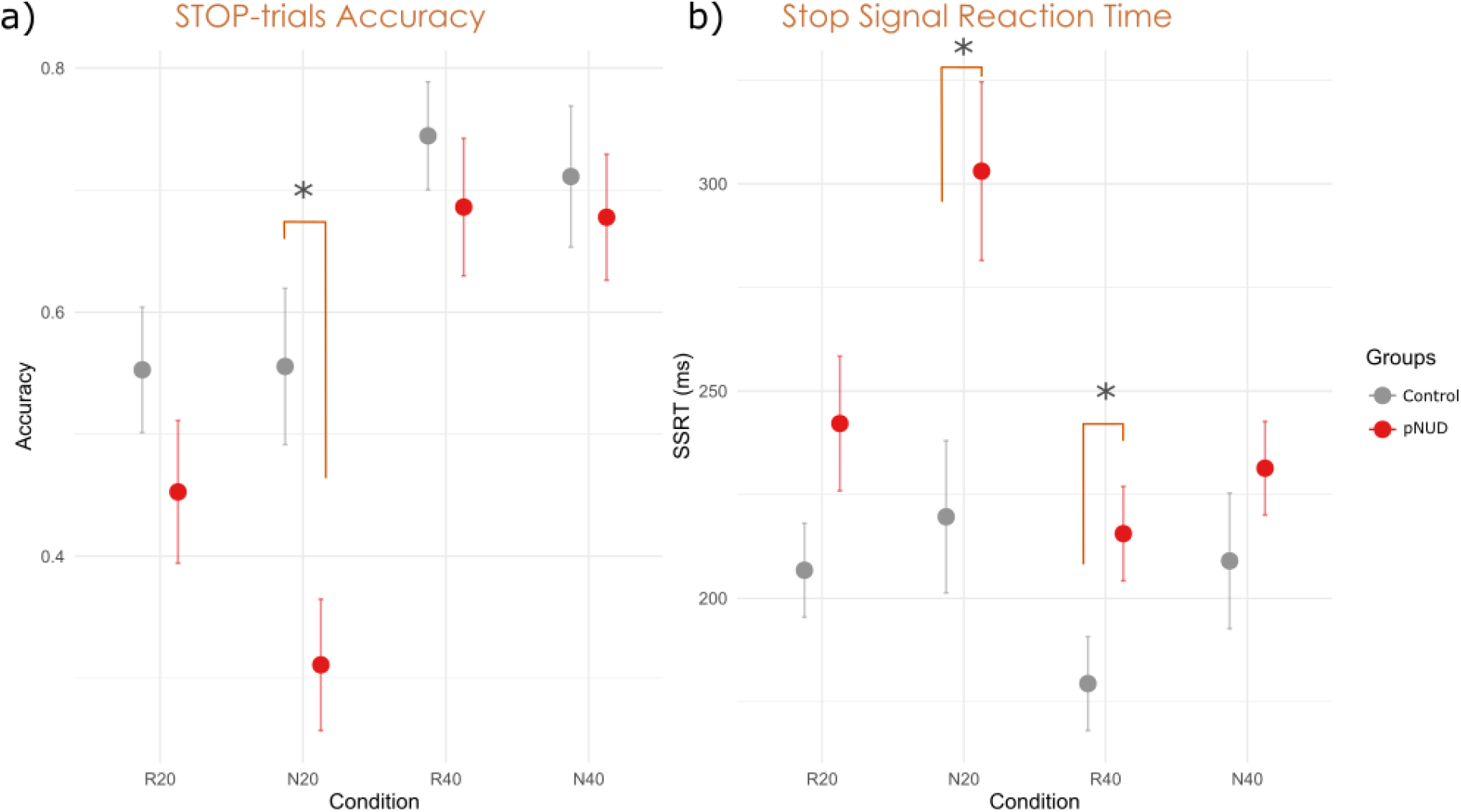
Inhibition accuracy measures for the people with a nicotine use disorder (pNUD) and control groups. **a)** *Stop trial accuracy scores*. Both groups show similar performance here, with no main effect group F(1,38) = 3.2, p = 0.08, η_p_^2^ = 0.078. There is a main effect of probability F(1,38) = 65.95, p < 0.0001*, ηp = 0.63, and group x probability interaction F(1,38) = 4.70 p = 0.036*, ηp = 0.11. There was a significant difference between N20 condition for control and pNUD group (t-test, p < 0.01). There were no significant main effect of reward (F(1,38) = 2.3, p = 0.14, η_p_^2^ = 0.058), no group and reward interaction (F(1,38) = 1.3, p = 0.1, ηp = 0.026) and no three way group, probability and reward interaction (F(1,38) = 2.98, p = 0.09, 0.073). **b)** The pNUD group have slower SSRTs (main effect of group: F(1,37) = 8.09, p < 0.01, η_p_^2^ = 0.18). There is a significant main effect of reward (F(1,37) = 12.96, p < 0.001, ηp = 0.26)* and probability (F(1,37) = 13.13, p < 0.001, η_p_^2^ = 0.26)*. There is a trend of a three way interaction between group, probability and reward (F(1,37) = 3.92, p = 0.055, ηp = 0.096). There were no significant two way interactions between group and reward (F(1,37) = 1.1, p = 0.3, η_p_^2^ = 0.03), reward and probability (F(1,37) = 0.8, p = 3.7, η_p_^2^ = 0.02) and group and probability (F(1,37) = 2.57, p = 0.12, ηp = 0.065). *\* = p < 0.05. pNUD = nicotine use disorder*.

#### pNUD have slower SSRTs compared to controls

Figure 3b shows SSRT estimates where pNUD had a slower SSRT compared to controls (main effect of group: F(1,37) = 8.09, p < 0.01, η_p_^2^ = 0.18). While pNUD had a similar stop-accuracy score, the slower SSRT suggest the pNUD group found inhibiting a response more effortful than controls. Independent t-tests indicated the pNUD group had a significantly slower SSRT for the N20 (p = 0.0054) and R40 (p = 0.034). There were no significant differences in the N40 (p = 0.25) and R20 (p = 0.097) conditions. We found a main effect of probability (F(1,37) = 13.13, p < 0.001, η_p_ ^2^ = 0.26) where the 40% condition had faster SSRTs. We also found a main effect of reward (F(1,37) = 12.96, p < 0.001, η_p_^2^ = 0.26) where reward trials had faster SSRTs. Lastly, there was no significant group and reward interaction (F(1,37) = 1.1, p = 0.3, η_p_^2^ = 0.03). Overall, we find that the pNUD group is more ‘impulsive’ in their responding based on their slower SSRTs.

### Brain fMRI Activity

#### Cue and Anticipation Epochs (preparatory control activity)

During the cue epoch here is more activity in the striatum for reward trials, compared to neutral (Figure 4a). Failed inhibitions have greater precentral and posterior medial frontal cortex (pmFC) activity (Figure 4b). The pNUD group exhibits more control related activity in the IFG, middle frontal gyrus (MFG) and superior frontal gyrus (SFG) compared to non-smokers, in reward trials which were successfully inhibited (Figure 4c). The pNUD group also has more activity in the pre and postcentral gyri prior to successful inhibitions in the reward trials. Lastly, there was more activity in the fusiform gyrus for the 40% > 20% probability contrast (figure not shown), which is consistent with the literature on the fusiform gyrus’ activation during the stop signal task, playing a role in correctly recognizing cues and their salience in inhibiting a response ^33–35^.

**Figure 4.**
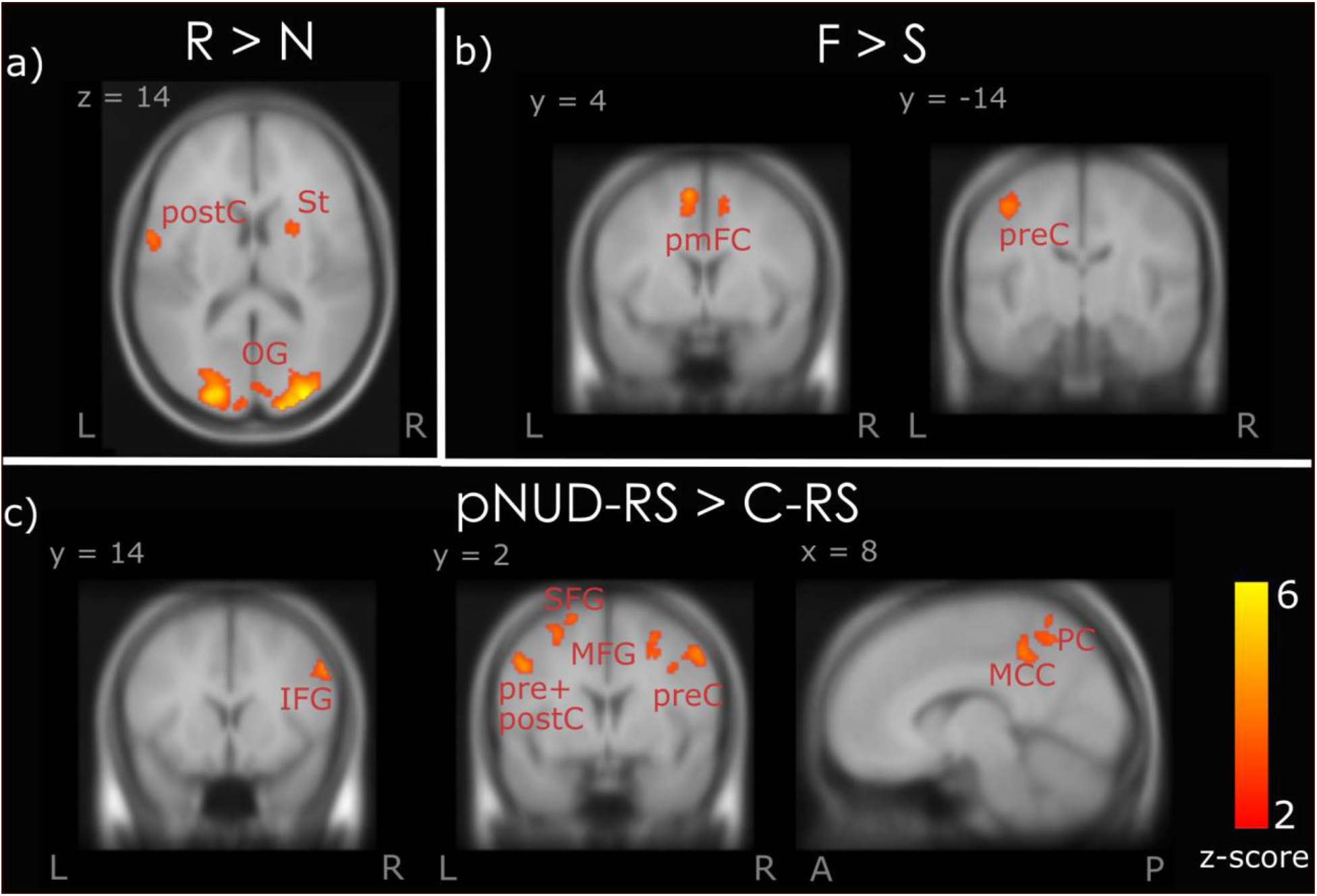
Activity during cue epoch. This is activity when participants are cued on the probability of an upcoming stop trial and whether it is a reward or a neutral trial. *Abbreviations: R = reward, N = neutral, F = Failed inhibition, S = successful inhibition, pNUD = people with a nicotine disorder group, C = control group. PC = precuneus, St = striatum, pmFC = posterior medial frontal cortex, IFG = inferior frontal gyrus, MFG = middle frontal gyrus, SFG = superior frontal gyrus, preC = precentral, postC = postcentral, MCC = mid cingulate cortex, OG = occipital gyrus. L = left, R = Right. A = anterior, P = posterior. *MFG + SFG activity in Figure 3c is pFWE = 0.055 (cluster-level)*.

There is greater striatal and insula activity at the anticipation epoch for reward > neutral trials (Figure 5a). Figure 5b shows activity in angular gyrus prior to successful inhibitions, consistent with previous literature ^9^. Failed inhibitions (Figure 5c) have more activity in the pre and postcentral gyri. There is also more activity in the medial cortical regions (anterior cingulate cortex; ACC, mid cingulate cortex; MCC and posterior medial frontal cortex; pmFC) associated with these failed inhibitions. The pNUD group in the anticipation epoch exhibits more control related activity in the IFG and MFG, as well as pre and postcentral activity (Figure 5d). Interestingly, pNUD in this epoch have more IFG activity even prior to failed inhibitions in the neutral trials (Figure 5e). See Table S1 and S2 in supplementary materials for a full list of brain regions activated in these epochs and their respective MNI coordinates and individual group activities).

**Figure 5.**
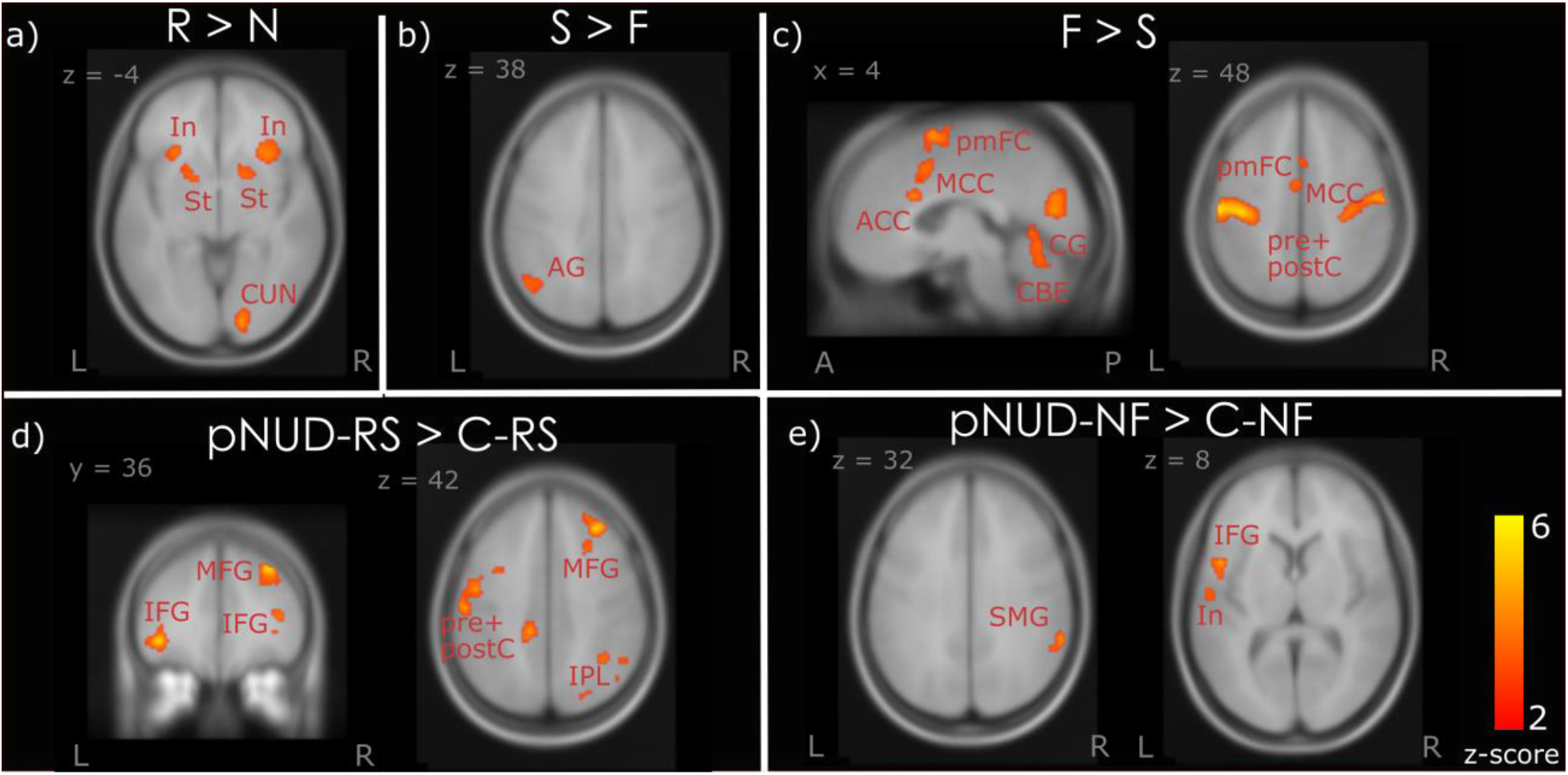
Brain activity during the anticipation epoch. This is activity immediately prior to the onset of the trial. *Abbreviations: R = reward, N = neutral, F = Failed inhibition, S = successful inhibition, pNUD = people with a nicotine disorder group, C = control group, RS = reward successful inhibition trials, NF = neutral failed inhibition trials In = insula, St = striatum, CUN = cuneus, ACC = anterior cingulate cortex, MCC = midcingulate cortex, pmFC = posterior medial frontal cortex, preC = precentral, postC = postcentral, IFG = inferior frontal cortex, SMG = supramarginal gyrus, MFG = middle frontal cortex, IPL = inferior parietal lobule, CBE = cerebellum. L = left, R = Right. A = anterior, P = posterior*.

#### Stop Trial Epoch (Reactive control activity)

Figure 6a shows the components of reactive control, where participants inhibit the prepotent response after seeing the stop signal. These successful inhibitions were associated with greater activity in control related regions (ACC, SFG, MFG), as well as the angular gyrus (Figure 6a). Successfully inhibiting reward trials engages more IFG, insula and striatum compared to successfully inhibiting neutral trials (Figure 6b). Non-smokers have greater ACC activation compared to smokers (Figure 6c), and this may suggest that non-smokers have more reactive control. See Table S3 in supplementary materials for a full list of brain regions activated in this epoch and their respective MNI coordinates and individual group activities.

**Figure 6.**
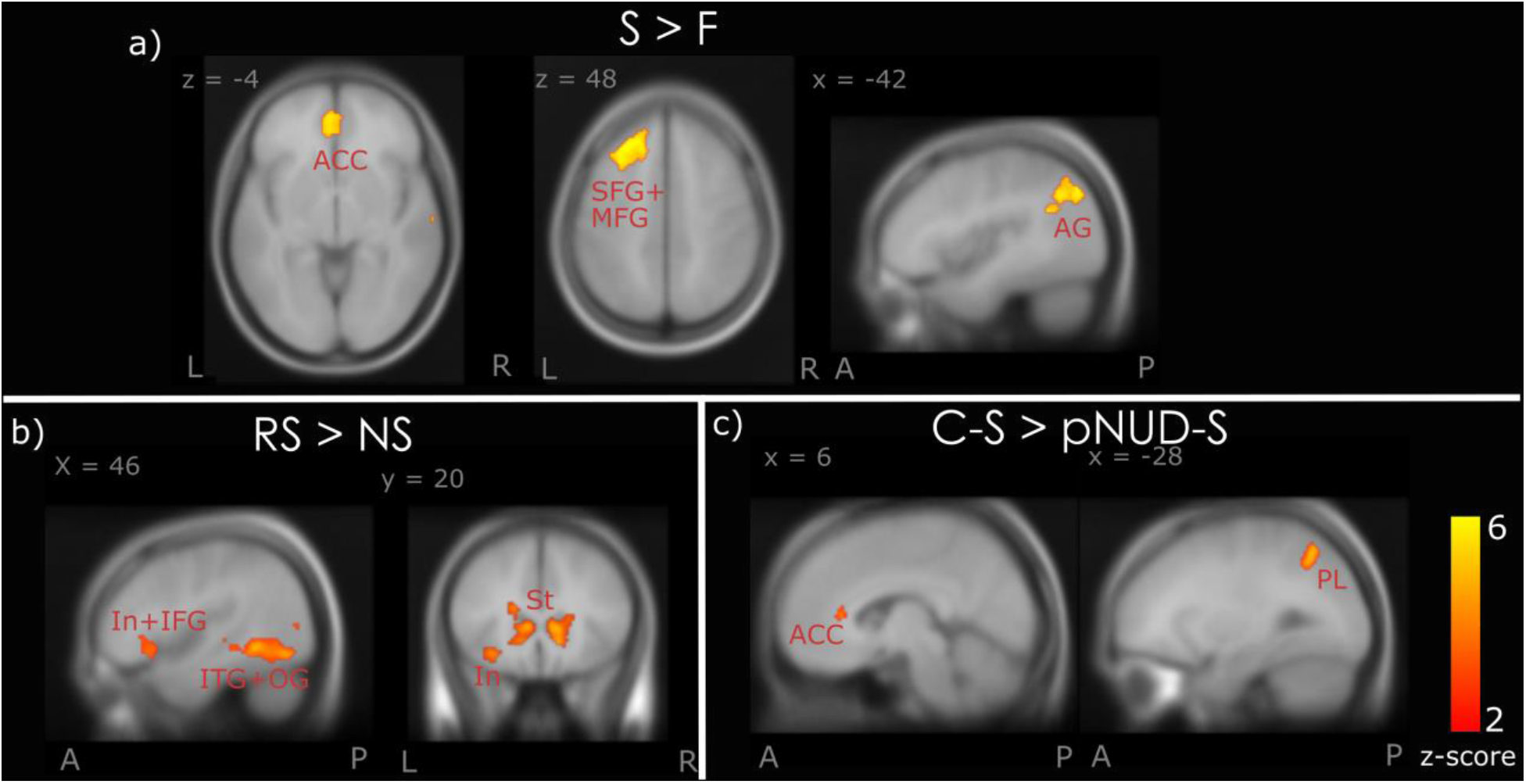
Brain activity at the stop-trial epoch. This is activity at the trial – when the stop signal is presented, and participants need to inhibit a response to make progress towards the larger later reward. *Abbreviations: R = reward, N = neutral, F = Failed inhibition, S = successful inhibition, RS = reward successful inhibition trials, NS = neutral successful inhibition trials, pNUD = people with a nicotine disorder group, C = control group. ACC = anterior cingulate cortex, SFG = superior frontal gyrus, MFG = middle frontal gyrus, In = insula, IFG = inferior frontal gyrus, St = striatum, AG = angular gyrus, PL = parietal lobule. L = left, R = Right. A = anterior, P = posterior. *The successful > failed inhibitions contrast here used threshold of p < 0.05 FWE corrected at whole brain level*

#### Feedback Epoch

Brain activity when processing feedback of “correct 0c” after successful inhibitions, contrasted with “miss 0c” after an incorrect response is shown in Figure 7a. Interestingly, there is striatal activity after successful inhibitions. Following successful inhibitions, there was also more activity in the ACC, SFG and MFG. In contrast, there was more insula and pmFC activity following failed inhibitions (Figure 7b). The pNUD group exhibited greater pmFC and SFG activity following failed inhibitions in neutral trials compared to controls (Figure 7c). In contrast, after successful inhibitions for reward trials, the pNUD group had greater activity in the medial cortical regions (cingulate gyrus, SFG, pmFC) and the IFG. See Table S4 in supplementary materials for a fill list of brain regions activated in this epoch and their respective MNI coordinates and individual group activities.

**Figure 7.**
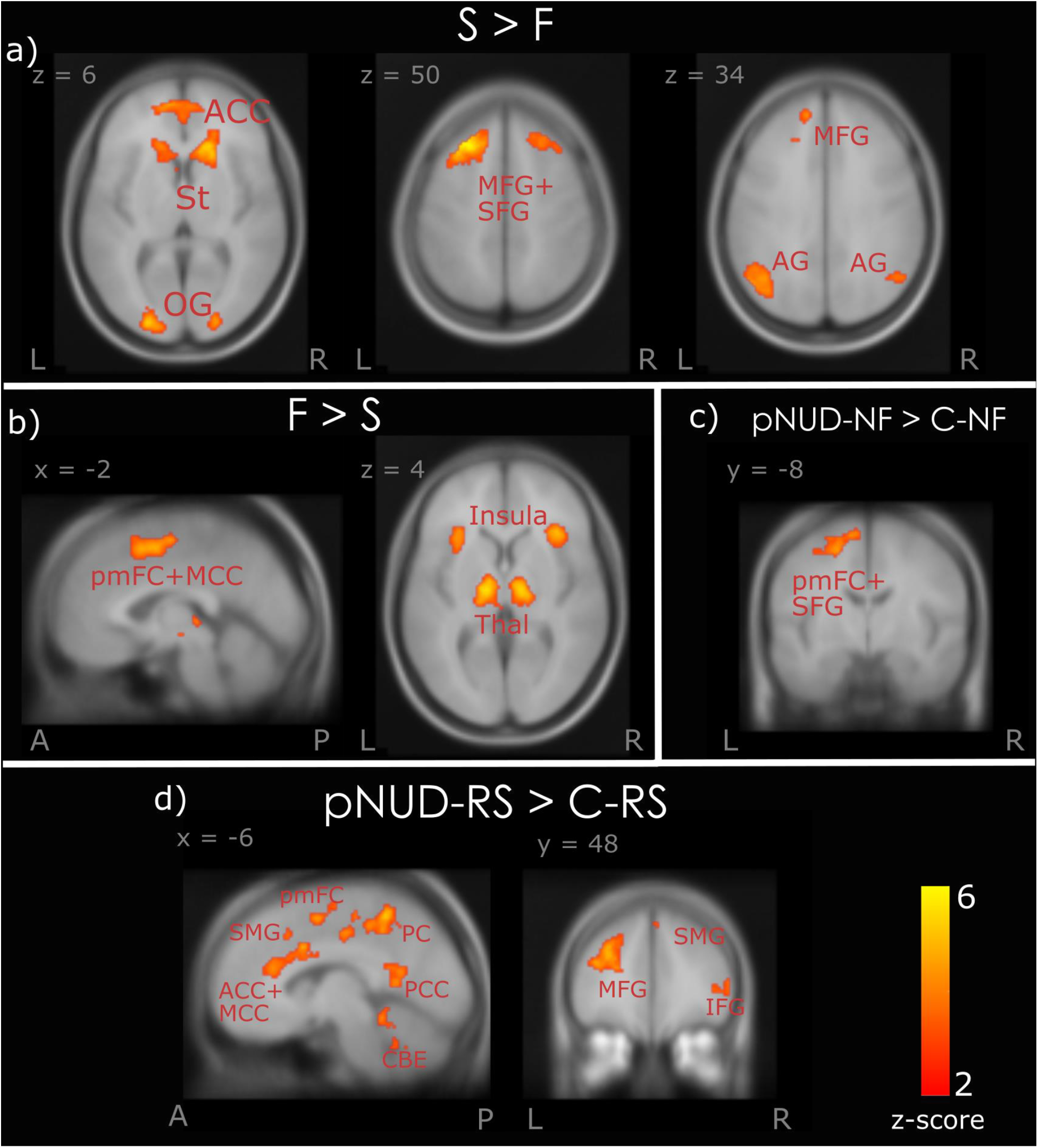
Brain activity at the feedback epoch. This is activity at the feedback epoch, where participants are given feedback of “correct 0c” for successful inhibitions and “incorrect 0c” for failed inhibitions, where the response was slower than 400ms or was the incorrect button press. *Abbreviations: R = reward, N = neutral, F = Failed inhibition, S = successful inhibition, RS = reward successful inhibition trials, NS = neutral successful inhibition trials, pNUD = people with a nicotine disorder group, C = control group. ACC = anterior cingulate cortex, SFG = superior frontal gyrus, MFG = middle frontal gyrus, In = insula, IFG = inferior frontal gyrus, MCC = midcingulate gyrus, PCC = posterior cingulate cortex, pmFC = posterior medial frontal cortex, OG = occipital gyrus, St = striatum, Thal = thalamus, CBE = cerebellum, SMG = superior medial gyrus, AG = angular gyrus. L = left, R = Right. A = anterior, P = posterior*.

#### Model-based SSRT brain activity correlations

The SSRT estimates of each participant across all conditions, between both groups, were used as covariates at the second level analysis (Figure 8). Across all epochs, we found that with slower SSRTs in the pNUD group, there is more activity within the PFC and parietal regions.

**Figure 8.**
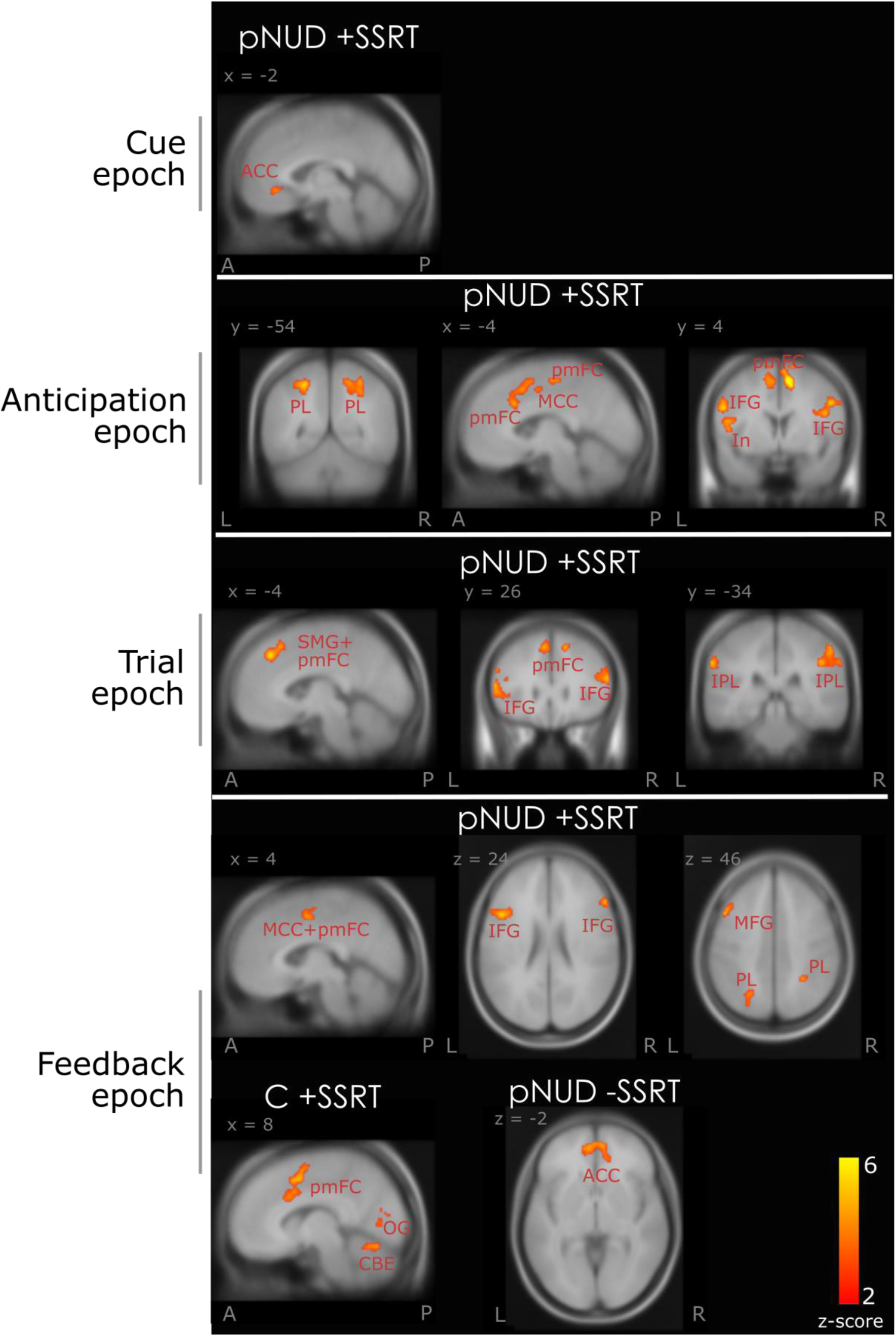
Brain activity correlating with SSRTs. pNUD+ SSRT = slower (increasing) SSRT correlating with the people with a nicotine disorder group. D-SSRT = faster (decreasing) SSRT correlating with the people with a nicotine disorder group. C+ SSRT = slower (increasing) SSRT correlating with the control group. *Abbreviations: ACC = anterior cingulate cortex, MCC = midcingulate cortex, pmFC = posterior medial frontal cortex, IFG = inferior frontal gyrus, SFG = superior frontal gyrus, MFG = middle frontal gyrus, IPL = inferior parietal lobe, PL = parietal lobule, CBE = cerebellum, OG = occipital gyrus*.

For the control group, we did not see SSRT brain correlations except for in the feedback epoch where a slower SSRT correlates with greater activity in the pmFC. See Table S5 in supplementary materials for a full list of brain regions activated in this epoch and their respective MNI coordinates.

## DISCUSSION

Here we investigated the underlying brain processes involved in exerting control over rewards in pNUD and healthy controls. Overall, we found that pNUD do not differ from controls in their stop-accuracy scores (Figure 3a). However, they had slower SSRTs (Figure 3b). The combination of these findings suggests that pNUD require more effort and/or have a greater difficulty inhibiting their prepotent ‘go-processes’. This is consistent with previous findings where slower SSRTs were found in pSUD, including to nicotine^12,13^ and a slower SSRT predicts a higher dependence to nicotine^36^. There was also a significant difference in the stop-accuracy score between groups specifically in the N20 condition, where the pNUD group had lower stop-accuracy scores. The difference here may be due to the task design where neutral and reward conditions are interleaved. Another speculative possibility is that the dependant group may prioritise their limited cognitive resources on inhibiting in other conditions, since these other conditions may either lead to a reward (for R20 and R40 conditions) or have a higher chance of being a stop trial (N40 condition). Overall, the pNUD group’s response inhibition performance was comparable to controls, but achieving parity required more cognitive effort.

The pNUD group did have greater activity in their cognitive control related prefrontal regions (IFG and MFG) prior to successful inhibitions for reward trials. This was during both, the cue and anticipation epochs. The increased activity might suggest the pNUD group engages in more preparatory control than healthy controls to achieve similar stop accuracies. Indeed, greater preparatory activity in prefrontal regions have previously been shown to aid in inhibitory processes^8,9^. Further, increased IFG activation for more difficult stops were previously reported^37^. The hypothesis that increased IFG activity in pNUD indicates more effort in stopping, is consistent with our SSRT results. This finding of increased preparatory control in the pNUD group is contrary to another recent finding, in people with a cocaine use disorder, found to have reduced PFC related preparatory contol^23^. One explanation for the contrasting results may be due to the differences in the severity of dependence, where our sample had close to moderate levels and therefore may have a greater inhibitory control capacity relative to people with a cocaine use disorder. A second possibility is our short 3-hour abstinence window, which may still produce some stimulant-related effects of nicotine and possibly cotinine (a metabolite of nicotine with a much longer half-life^38^) on cognition and may therefore facilitate this “adaptive” pattern of behavior where reduced reactive control is compensated for by increased preparatory control. Interestingly, the pNUD group also exhibits greater activity in the pre and postcentral gyri, prior to successful inhibitions in reward trials. The increased precentral motor-related activity may be inhibited by the increased prefrontal control activity, contributing to the successful stop. Another possibility is that the precentral activation may be playing a role in motor inhibition as supported by findings from Li, Huang, Constable, & Sinha (2006), where precentral activation correlated with smaller SSRTs (or efficient inhibitions).

The pNUD in the anticipation epoch had more IFG activity prior to failed inhibitions in neutral trials (Figure 5e). Increased IFG activity prior to failed inhibitions observed here suggests that greater preparatory IFG activity does not guarantee a successful inhibition and may also need engagement from other control related prefrontal regions, such as MFG, as is the case prior to the successful inhibitions (Figure 5d). The finding that IFG may not guarantee a successful inhibition supports previous studies which find that IFG may play a non-specific role in response inhibition, for example, encoding other aspects of the task, such as attention, uncertainty and salience detection^40^. Overall, the pNUD group engages more preparatory control activity prior to successful inhibitions in reward trials, aiding them in achieving similar stop accuracy scores to the controls, albeit with slower SSRTs. During the trial epoch, when the stop signal is detected, we see components of reactive control over reward. Successful inhibitions, across both groups, engaged more prefrontal control regions (MFG, SFG) and the ACC. These regions may aid in successfully inhibiting the go-processes and therefore contribute to the successful stop. The ACC has been suggested to play a more complicated and non-specific role in the stop signal task^39^, however, one of its key roles has been implicated in inhibiting responses^41^. We therefore interpret ACC activity here as playing a role in facilitating response inhibition. Interestingly, the control group had greater ACC activity compared to the pNUD group during successful inhibitions in reward trials, suggesting that while the pNUD group may have an increased preparatory control, the control group have greater reactive control. The combination of these findings suggests that greater preparatory control (with increased IFG and MFG activity) reduces the need for high levels of reactive control to inhibit a prepotent response; as is the case for the dependent group. The control group, on the other hand, show less preparatory control but increased reactive control (with increased ACC activity) and can still reliably inhibit their responses. The control group may therefore find it less difficult to inhibit their responses and may not require the upregulated preparatory control for a successful stop. Our SSRT results showing the control group have shorter SSRTs is consistent with this interpretation in that they may find it less difficult to inhibit responses and therefore may rely less on increased preparatory control due to their higher levels of reactive control. Overall, both groups have similar inhibition accuracy, but achieve this through different processes. At the feedback epoch, there was increased striatal activation following successful inhibitions when contrasted with failed inhibitions, in both groups. Given the striatum’s role in reward processing^16^, striatal activity here is consistent with participants anticipating the larger $20 reward and this may play a role in motivating further response inhibitions to obtain the larger later reward. Further, the feedback of “correct 0c” contrasted with “miss 0c” may also exhibit a component of intrinsic reward processing for correctly performing the task. Contrary to our hypothesis, we did not see striatal activity during impulsive responses (for the smaller 20c reward), as previous studies have found^17^. Instead, the striatum was engaged during successful inhibitions in reward trials over neutral (Figure 6b). This may be due to the participants’ goal of attaining the larger later reward as compared to the smaller sooner, where progress towards the larger later reward engaged striatal anticipatory activity. Therefore, the small 20c “reward” may be considered neutral or even punishing by the participants given the context of the trials, and the overall goal of attaining the larger later reward.

Following successful inhibitions (in the feedback epoch), there was also more activity in the ACC, SFG and MFG. These brain regions have previously been implicated in processing feedback, including positive feedback^42,43^. Interestingly, the pNUD group had greater activity in the medial cortical regions and the IFG, following successful inhibitions in reward trials, compared to controls. Feedback processing and associated activity in these medial cortical regions have previously been shown to increase task performance ^43^. One interpretation may be that increased activation in these regions for the pNUD group aids in improving stop accuracy scores to match those of the control group. Decreases in inhibitory control may lead to different compensatory neural strategies by the pNUD group and increasing feedback processing during successful inhibitions may therefore be one such compensatory neural strategy.

The model-based SSRT brain correlations showed that the pNUD group has more cognitive control related activity correlating with slow SSRTs, across all epochs. On the contrary, the control group shows more control related SFG and precentral activity to correlate with fast SSRTs, although only at a relaxed statistical threshold (SFG; p = 0.066 FWE (cluster-level) and precentral; p = 0.057 FWE (cluster-level)). Overall, one may expect greater engagement of these control related regions to faster SSRTs, hence enabling more efficient stopping, as found by Galván, Poldrack, Baker, McGlennen, & London (2011), where both smokers and non-smokers had greater activity in control related regions that correlated with faster SSRT. However, Galván and colleagues (2011) did not find SSRT differences between groups. Other studies correlating SSRT scores with fMRI data have found that faster SSRTs correlate more strongly with control related regions (medial cortical regions, SFG) and motor regions such as the pre-supplementary motor area and precentral^39,45,46^. These were all investigated in healthy non-smoker participants. Consistent with these studies, we found more precentral and PFC activity for the control group. However, for the pNUD group it appears that slower SSRT engages more control related regions. One interpretation to bring together these SSRT results across groups may be that the SSRTs reflect the cognitive effort required for stopping, or difficulty in inhibiting. For the control group, more effort is required for fast inhibitions, hence greater control related SFG activity. In contrast, the pNUD group appears to find the slower SSRTs more effortful, consistent with greater control related PFC activity.

In sum, we find that both control and pNUD participants exhibit similar stop accuracy scores. However, the brain processes exhibited to achieve this are different between the two groups. The pNUD group shows greater inhibitory control related activity prior to successful inhibitions over reward, whereas, the control group exhibit greater reactive control with greater ACC activity during the inhibition epoch. Collectively, our results shed light on some of the brain processes involved in successfully exhibiting control over immediate small rewards in favor of greater delayed gratification in non-smokers and pNUD.

## Acknowledgments

The project was funded by the NHMRC project grant 1050766 to Robert Hester. Shivam Kalhan was supported by the Velma Stanley PhD Scholarship and by the Australian Government Research Training Program Scholarship, provided by the Australian Commonwealth Government and the University of Melbourne.

## Author Contributions

All authors contributed to this paper. E.C collected the data and performed preliminary analysis. R.H designed the experiment and wrote the paper. M.I.G wrote the paper. S.K analyzed data and wrote the paper.

## Conflict of Interest

None.

## Data Availability Statement

The data that support the findings of this study are available on request from the corresponding author. The data are not publicly available due to privacy or ethical restrictions.

## Notes

### Competing Interest Statement

The authors have declared no competing interest.

