## Supplementary Materials for "People with nicotine use-disorder exhibit more prefrontal activity during preparatory control but reduced anterior cingulate activity during reactive control"

### Supplementary Material

#### *Supplementary Methods*

*Apparatus*

The behavioral task was presented using E-Prime® 2.0 software (Psychology Software Tools, Pittsburgh, PA). The MRI scanner used was a 32-channel head coil Siemens Tim Trio 3T MRI (Germany). Behavioral responses in scanner were recorded using a two-bottom response box (Fibre-Optic response pad, Current Designs, Philadelphia, PA, USA). Functional MRI data was analyzed using SPM12 (https://www.fil.ion.ucl.ac.uk/spm/software/spm12/). Behavioral analysis was done using R (https://www.r-project.org/) and SPSS (ibm.com/au-en/analytics/spss-statistics-software). Behavioral data visualizations were done with the “ggplot2” package in R software (https://ggplot2.tidyverse.org/reference/ggplot.html).

*Procedure*

Two sessions were conducted on two separate days. The first session (approx. 30 minutes), was held at the University of Melbourne to screen participants and familiarize them with the behavioral task. The second session (approx. 45 minutes), administered a longer version of the behavioral task in the MRI scanner. The scanner was based at Swinburne University (Melbourne, Australia). All participants were reimbursed $20 per hour, plus additional payments depending on task performance for both sessions.

Before the start of the first session, participants completed the consent form, demographic and screening questionnaires. Participants were briefed on the task with the experimenter providing verbal instructions during practice of the task, particularly focusing on the different monetary reward contingences. At the beginning of the second session, participants were briefed again and performed a short version of the behavioral task. In ensuring that participants were aware that the money they may receive is real, they were informed that they would be reimbursed for their time spent at the MRI facility in addition to receiving the monetary reward based on their task performance.

*MRI Sequences*

Functional images were acquired using T2* echo-planar imaging (EPI) sequences, with six runs in total. These were acquired using the following parameters: repetition time (TR) = 2s; echo time (TE) = 36 ms: flip angle (FA) = 90$^{o}$; 192mm field of view (FOV); and 38 contiguous slices of 4mm slice thickness. The first three volumes of each run were discarded prior to data analysis to account for transients in the magnetic field of scanner. Following functional imaging, structural images were acquired using an MPRAGE T1-weighted with the following sequences: TR = 1900ms; TE = 2.3ms, FA = 90, slice thickness = .90mm; in-plane resolution = 1 x 1mm$^2$.

*Stop Signal Reaction Time Estimation*

The SSRTs were estimated with the horse-race model, using the integration method[1]. The estimation steps were followed as outlined in Eagle et al., (2008) and were estimated per participant, and per condition. Therefore, each participant had an SSRT value estimate for each of the four conditions (R20, N20, R40, N40). The integration method used assumes that SSRTs are constant, however, this assumption does not affect the reliability of SSRT estimates [3–5]. It has been suggested that 50 or more stop trials can produce reliable SSRT estimates, however, our task had fewer stop-trials than this per condition (18 for each reward condition and 9 for each neutral condition; total = 54)[6]. The fewer number of stop-trials per condition is therefore a possible limitation in our task. Additionally, our paradigm consists of one fixed SSD as supposed to a variable SSD with a tracking procedure. This presents with another limitation where participants may wait for the stop signal to appear or be able to predict when it will occur within a trial [6]. This may possibly bias our SSRT estimates.

***Supplementary Results***

*Both groups use a less cautious approach in responding for reward and 20% probability go-trials*

For both groups, the 40% probability conditions have a smaller proportion of go-trial responses under the 400ms threshold (main effect probability: $F(1,38) = 40$, $p < 0.0001$, $\eta_p^2 = 0.51$) (Figure 4a) and a slower reaction time (main effect probability: $F(1,38) = 44.28$, $p < 0.00001$, $\eta_p^2 = 0.54$) (Figure 4b), compared to the 20% probability conditions. This may be indicative of lesser difficulty (and a more cautious strategy being used) in inhibiting a response at the 40% probability conditions, compared to the 20% probability condition. In contrast, the reward trials have a greater proportion of go-trial responses under 400ms (main effect of reward: $F(1,38) = 11.6$, $p < 0.05$, $\eta_p^2 = 0.23$) and a faster reaction time (main effect of reward: $F(1,38) = 24.74$, $p < 0.0001$, $\eta_p^2 = 0.4$), compared to the neutral conditions. This is indicative of a less cautious approach when a quick response may lead to the monetary reward. Lastly, there was no significant main effect of group or interactions from the go-trial accuracy and reaction times (see Figure 4 legend for the statistics). Overall, we find that both groups use a more cautious approach to responding in go-trials for the 40% probability condition and a less cautious approach for the reward trials, where a quick response may lead to the small 20¢ reward.

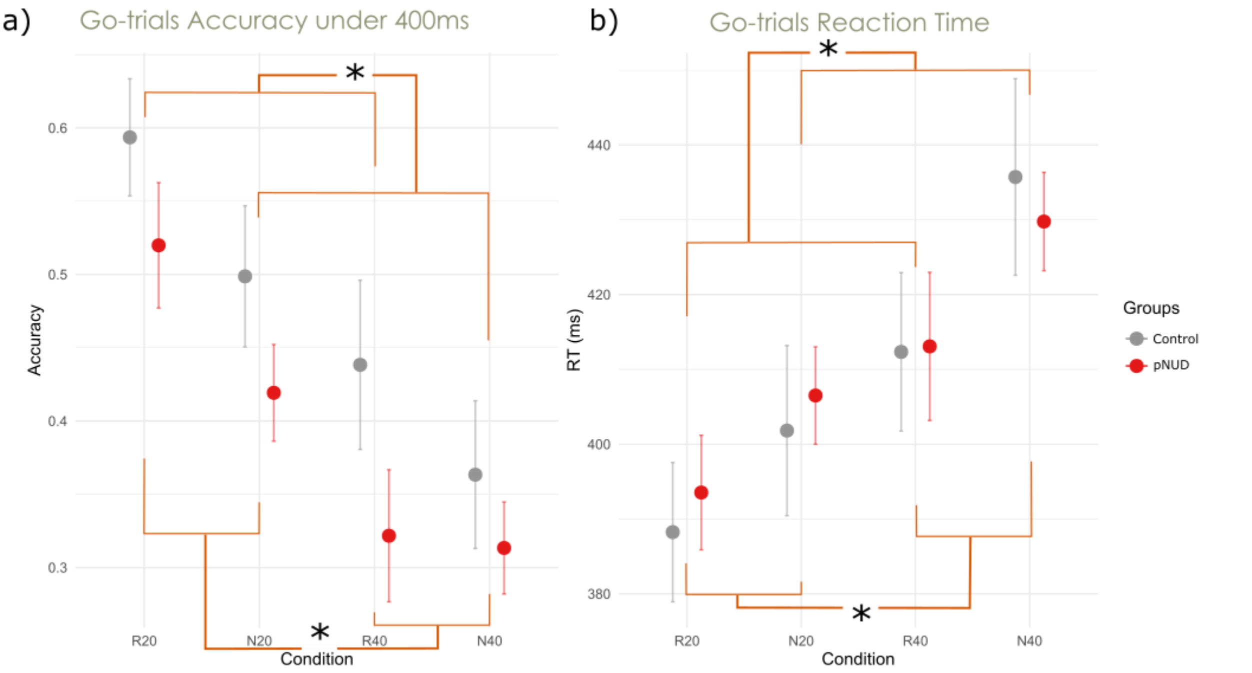


***Figure S1.** Behavioural measures for go-trials.* ***a)*** Go trials accuracy scores for responses under 400ms. There was no main effect of group ($F(1,38) = 2.32$, $p = 0.14$, $\eta_p^2 = 0.06$). There was a significant main effect of reward ($F(1,38) = 11.6$, $p < 0.05$, $\eta_p^2 = 0.23$)* and probability ($F(1,38) = 40$, $p < 0.0001$, $\eta_p^2 = 0.51$)*. There was no significant group x reward interaction ($F(1,38) = 0.56$, $p = 0.5$, .014), group x probability interaction ($F(1,38) = 0.02$, $p = 0.9$, $\eta_p^2 = 0.001$), reward x probability interaction ($F(1,38) = 4$, $p = 0.053$, $\eta_p^2 = 0.095$) and group x probability x reward interaction ($F(1,38) = 1.67$, $p = 0.2$, $\eta_p^2 = 0.042$). ***b)*** Go trials reaction times. There was no significant main effect of group ($F(1,38) = 0.01$, $p = 0.92$, $\eta_p^2 = 0.0001$). There was a significant main effect of reward ($F(1,38) = 24.74$, $p < 0.0001$, $\eta_p^2 = 0.4$)* and probability ($F(1,38) = 44.28$, $p < 0.00001$, $\eta_p^2 = 0.54$)*. There were no significant interactions; group x reward ($F(1,38) = 0.3$, $p = 0.6$, $\eta_p^2 = 0.008$), group x probability ($F(1,38) = 1$, $p = 0.3$, $\eta_p^2 = 0.026$), reward x probability ($F(1,38) = 1.8$, $p = 0.2$, $\eta_p^2 = 0.045$) and group x reward x probability ($F(1,38) = 0.4$, $p = 0.6$, $\eta_p^2$ = 009). *pNUD = people with a nicotine use disorder.*

Table S1a: Reward > Neutral (R > N) contrast at cue epoch

| **Brain Region** | **Side** | **X** | **Y** | **Z** |
|---|---|---|---|---|
| Inferior Occipital Gyrus | L | -22 | -90 | -10 |
| Lingual Gyrus | R | 26 | -86 | -8 |
| Middle Occipital Gyrus | R | 30 | -92 | 6 |
| Thalamus | L | -22 | -28 | -4 |
| Postcentral Gyrus | R | 52 | -16 | 36 |
| Precentral Gyrus | R | 44 | -6 | 44 |
| Postcentral Gyrus | L | -58 | 2 | 14 |

| | | | | |
|---|---|---|---|---|
| Postcentral Gyrus | L | -60 | -2 | 28 |
| Postcentral Gyrus | L | -62 | -10 | 22 |
| Striatum (Caudate Nucleus) | R | 12 | 14 | -4 |
| Thalamus | R | 10 | -4 | -2 |

Table S1b: Failed > Successful (F > S) inhibitions at cue epoch

| Brain Region | Side | X | Y | Z |
|---|---|---|---|---|
| Posterior-Medial Frontal | L | -8 | 2 | 58 |
| Posterior-Medial Frontal | L | -10 | 2 | 48 |
| Posterior-Medial Frontal | R | 12 | 0 | 54 |
| Precentral Gyrus | L | -40 | -16 | 50 |
| Precentral Gyrus | L | -28 | -22 | 54 |
| Precentral Gyrus | L | -26 | -28 | 62 |

Table S1c: Successful inhibitions at reward trials, pNUD group > control group (pNUD-RS > C-RS) at cue epoch

| Brain Region | Side | X | Y | Z |
|---|---|---|---|---|
| Precentral Gyrus | L | -50 | 4 | 38 |
| Postcentral Gyrus | L | -56 | -6 | 42 |
| Precentral Gyrus | R | 54 | 4 | 42 |
| Inferior Frontal Gyrus (p. Opercularis) | R | 54 | 14 | 36 |
| Precuneus | L | -6 | -48 | 50 |
| Mid Cingulate Gyrus | R | 8 | -40 | 46 |
| Precuneus | R | 6 | -58 | 54 |
| Middle Frontal Gyrus | L | -28 | 4 | 54 |
| Superior Frontal Gyrus | L | -20 | 2 | 64 |
| Posterior-Medial Frontal | L | -16 | -6 | 62 |

Table S2a: Reward > Neutral (R > N) contrast at anticipation epoch

| Brain Region | Side | X | Y | Z |
|---|---|---|---|---|
| Cuneus | R | 18 | -98 | 10 |
| Lingual Gyrus | R | 16 | -90 | -6 |
| Cerebellum | R | 10 | -74 | -16 |
| Insula Lobe | R | 30 | 28 | 8 |
| Striatum (Caudate Nucleus) | R | 12 | 8 | -2 |
| Striatum (Putamen) | L | -20 | 10 | -2 |

| | | | | |
|---|---|---|---|---|
| Thalamus | R | 2 | -4 | 6 |
| Insula Lobe | L | -28 | 22 | -6 |
| Insula Lobe | L | -28 | 28 | 6 |

Table S2b: Successful > Failed (S > F) inhibitions contrast at anticipation epoch

| Brain Region | Side | X | Y | Z |
|---|---|---|---|---|
| Angular Gyrus | L | -44 | -62 | 46 |
| Angular Gyrus | L | -52 | -60 | 36 |

Table S2c: Failed > Successful (F > S) inhibitions contrast at anticipation epoch

| Brain Region | Side | X | Y | Z |
|---|---|---|---|---|
| Postcentral Gyrus | L | -54 | -20 | 46 |
| Rolandic Operculum (Insula) | L | -42 | -6 | 12 |
| Rolandic Operculum (Insula) | L | -52 | 4 | 14 |
| Thalamus | L | -14 | -22 | 4 |
| Thalamus | L | -12 | -12 | 12 |
| Calcarine Gyrus | L | -10 | -78 | 12 |
| Calcarine Gyrus | R | 14 | -74 | 14 |
| Cerebellum | L | -8 | -56 | -8 |
| Anterior Cingulate Cortex | R | 6 | 20 | 24 |
| Mid-Cingulate Cortex | L | -8 | 12 | 36 |
| Mid-Cingulate Cortex | R | 8 | 12 | 34 |
| Postcentral Gyrus | R | 56 | -14 | 48 |
| Supramarginal Gyrus | R | 56 | -16 | 26 |
| Precentral | R | 34 | -20 | 44 |
| Posterior-Medial Frontal | R | 6 | 8 | 66 |
| Posterior-Medial Frontal | L | 0 | -4 | 56 |
| Mid-Cingulate Cortex | L | -2 | -4 | 48 |
| Rolandic Operculum (Insula) | R | 42 | -2 | 16 |

Table S2d: Successful inhibitions at reward trials, pNUD group > control group (pNUD-RS > C-RS) at anticipation epoch

| Brain Region | Side | X | Y | Z |
|---|---|---|---|---|
| Inferior Frontal Gyrus (p. Orbitalis) | L | -40 | 36 | -4 |
| Inferior Frontal Gyrus (p. Triangularis) | L | -54 | 26 | 8 |
| Inferior Frontal Gyrus (p. Orbitalis) | L | -40 | 28 | -12 |
| Middle Frontal Gyrus | R | 32 | 36 | 44 |
| Middle Frontal Gyrus | R | 28 | 24 | 44 |

| | | | | |
|---|---|---|---|---|
| Inferior Parietal Lobule | R | 46 | -52 | 52 |
| Superior Parietal Lobule | R | 16 | -56 | 66 |
| Superior Parietal Lobule | R | 20 | -62 | 62 |
| SupraMarginal Gyrus | L | -62 | -36 | 30 |
| Middle Temporal Gyrus | L | -52 | -26 | -2 |
| Rolandic Operculum (Insula) | L | -60 | -8 | 8 |
| Middle Temporal Gyrus | R | 56 | -40 | 2 |
| Middle Temporal Gyrus | R | 54 | -38 | -6 |
| Middle Temporal Gyrus | R | 60 | -28 | -10 |
| Calcarine Gyrus | R | 10 | -62 | 18 |
| Precuneus | L | -8 | -52 | 8 |
| Precentral Gyrus | L | -48 | -6 | 48 |
| Postcentral Gyrus | R | -56 | -16 | 44 |
| Precentral Gyrus | R | -50 | -2 | 40 |
| Middle Frontal Gyrus | R | 40 | 38 | 12 |
| Middle Frontal Gyrus | R | 38 | 42 | 0 |
| Middle Frontal Gyrus | R | 46 | 46 | 8 |
| Superior Temporal Gyrus | R | 56 | -4 | -6 |
| Temporal Pole | R | 52 | 6 | -14 |
| Middle Temporal Gyrus | R | 58 | -58 | 14 |
| SupraMarginal Gyrus | R | 60 | -40 | 34 |
| Superior Temporal Gyrus | R | 58 | -52 | 20 |
| Mid-Cingulate Gyrus | L | -12 | -34 | 44 |
| Precentral Gyrus | L | -44 | 0 | 28 |
| Middle Frontal Gyrus | L | -28 | 6 | 46 |

Table S2e: Failed inhibitions at neutral trials, pNUD group > control group (pNUD-NF > C-NF) at anticipation epoch

| | | | | |
|---|---|---|---|---|
| Inferior Frontal Gyrus (p. Opercularis) | L | -48 | 8 | 8 |
| Postcentral Gyrus | L | -50 | -4 | 16 |
| Rolandic Operculum (Insula) | L | -50 | 0 | 8 |
| SupraMarginal Gyrus | R | 60 | -42 | 32 |
| Superior Temporal Gyrus | R | 58 | -50 | 20 |
| Middle Temporal Gyrus | R | 48 | -52 | 14 |

Table S3a: Successful > Failed (S > F) inhibitions contrast at trial epoch

| **Brain Region** | **Side** | **X** | **Y** | **Z** |
|---|---|---|---|---|
| Superior Frontal Gyrus | L | -16 | 28 | 52 |
| Middle Frontal Gyrus | L | -26 | 30 | 48 |
| Superior Frontal Gyrus | L | -14 | 40 | 46 |
| Precuneus | L | -10 | -54 | 42 |

| | | | | |
|---|---|---|---|---|
| Posterior cingulate cortex | L | -6 | -52 | 24 |
| Posterior cingulate cortex | R | 2 | -46 | 30 |
| Superior Temporal Gyrus | R | 64 | -24 | 2 |
| Angular Gyrus | L | -44 | -72 | 38 |
| Angular Gyrus | L | -52 | -62 | 34 |
| Angular Gyrus | L | -42 | -60 | 36 |
| Mid Orbital Gyrus | L | -4 | 50 | -6 |
| Anterior Cingulate Cortex | L | -4 | 42 | -4 |
| Superior Medial Gyrus | L | -4 | 56 | 10 |
| Superior Medial Gyrus | L | 14 | 54 | 2 |
| Middle Temporal Gyrus | L | -62 | -12 | -2 |
| Angular Gyrus | R | 54 | -62 | 30 |
| Superior Frontal Gyrus | R | 20 | 36 | 46 |
| Superior Frontal Gyrus | R | 14 | 50 | 38 |

Table S3b: Reward successful > Neutral successful (RS > NS) inhibitions contrast at trial epoch

| **Brain Region** | **Side** | **X** | **Y** | **Z** |
|---|---|---|---|---|
| Thalamus | R | 2 | -20 | 12 |
| Thalamus | R | 2 | -6 | 8 |
| Cerebellum | L | -8 | -80 | -22 |
| Thalamus | L | -28 | -30 | -2 |
| Occipital Gyrus/Lingula Gyrus | R | 18 | -76 | -12 |
| Insula Lobe | R | 10 | -72 | 56 |
| Insula Lobe/IFG (p. orbital) | R | 44 | 22 | -10 |
| Insula Lobe | R | 38 | 14 | -10 |
| Precuneus | R | 10 | -72 | 56 |
| Cuneus | R | 8 | -80 | 42 |
| Precuneus | L | -6 | -72 | 52 |
| Striatum | L | -16 | 12 | 20 |
| Striatum | R | 12 | 22 | 4 |

Table S3c: successful inhibitions, control group > pNUD group (C-S > pNUD-S) contrast at trial epoch

| **Brain Region** | **Side** | **X** | **Y** | **Z** |
|---|---|---|---|---|
| Middle Temporal Gyrus | L | -50 | -52 | -2 |
| Inferior Temporal Gyrus | L | -48 | -56 | -10 |
| Superior Parietal Lobule | L | -28 | -62 | 50 |
| Inferior Parietal Lobule | L | -30 | -58 | 42 |
| Superior Parietal Lobule | L | -22 | -54 | 52 |
| Inferior Temporal Gyrus | R | 48 | -70 | -10 |

| | | | | |
|---|---|---|---|---|
| Inferior Temporal Gyrus | R | 50 | -56 | -8 |
| Middle Temporal Gyrus | R | 54 | -60 | 2 |
| Anterior Cingulate Cortex | R | 10 | 30 | 8 |
| Anterior Cingulate Cortex | R | 16 | 38 | 8 |
| Superior Temporal Gyrus | R | 50 | -18 | -4 |
| Amygdala | R | 30 | 2 | -18 |

Table S4a: Successful > Failed (S > F) inhibitions contrast at feedback epoch

| **Brain Region** | **Side** | **X** | **Y** | **Z** |
|---|---|---|---|---|
| Striatum (Caudate Nucleus) | L | -12 | 20 | 0 |
| Anterior Cingulate Cortex | L | -4 | 48 | -4 |
| Superior Frontal Gyrus | L | -16 | 30 | 42 |
| Middle Frontal Gyrus | L | -22 | 28 | 48 |
| Superior Frontal Gyrus | L | -18 | 40 | 44 |
| Middle Occipital Gyrus | L | -22 | -92 | 4 |
| Cerebellum | L | -22 | -82 | -20 |
| Inferior Occipital Gyrus | L | -18 | -98 | -8 |
| Angular Gyrus | L | -40 | -58 | 28 |
| Angular Gyrus | L | -40 | -68 | 40 |
| Angular Gyrus | L | -48 | -66 | 38 |
| Superior Frontal Gyrus | R | 22 | 32 | 56 |
| Middle Frontal Gyrus | R | 34 | 28 | 52 |
| Superior Frontal Gyrus | R | 22 | 36 | 48 |
| Lingual Gyrus | R | 16 | -86 | -6 |
| Lingual Gyrus | R | 24 | -86 | -10 |
| Superior Occipital Gyrus | R | 22 | -92 | 4 |
| Angular Gyrus | R | 42 | -72 | 40 |
| Angular Gyrus | R | 50 | -60 | 36 |

Table S4b: Failed > Successful (F > S) inhibitions contrast at feedback epoch

| **Brain Region** | **Side** | **X** | **Y** | **Z** |
|---|---|---|---|---|
| Thalamus | L | -12 | -14 | 6 |
| Thalamus | R | 14 | -14 | 8 |
| Thalamus | R | 8 | -8 | -2 |
| Posterior-Medial Frontal | R | 4 | 14 | 48 |
| Mid-Cingulate Cortex | R | 8 | 14 | 34 |
| Posterior-Medial Frontal | L | 0 | 4 | 52 |
| Insula Lobe | L | -30 | 22 | -4 |
| Insula Lobe | L | -32 | 20 | 6 |
| Insula Lobe | R | 36 | 26 | 4 |

| | | | | |
|---|---|---|---|---|
| Insula Lobe | R | 40 | 22 | -4 |
| Cerebellum | R | 18 | -50 | -20 |
| Cerebellum | R | 22 | -60 | -24 |
| Cerebellum | R | 32 | -46 | -28 |

Table S4c: Failed inhibitions at neutral trials, pNUD group > control group (pNUD-NF > C-NF) contrast at feedback epoch

| Brain Region | Side | X | Y | Z |
|---|---|---|---|---|
| Superior Frontal Gyrus | L | -22 | -10 | 54 |
| Posterior-Medial Frontal | L | -10 | -8 | 64 |
| Middle Frontal Gyrus | R | 38 | -8 | 52 |
| Middle Temporal Gyrus | L | -58 | -50 | 10 |
| Middle Temporal Gyrus | L | -42 | -54 | 14 |
| Middle Temporal Gyrus | L | -50 | -52 | 10 |

Table S4d: Successful inhibitions at reward trials, pNUD group > control group (pNUD-RS > C-RS) at feedback epoch

| Brain Region | Side | X | Y | Z |
|---|---|---|---|---|
| Paracentral Lobule | R | 12 | -36 | 54 |
| Precuneus | L | -10 | -44 | 48 |
| Precuneus | L | -6 | -46 | 56 |
| Superior Frontal Gyrus | L | -12 | 6 | 54 |
| Posterior-Medial Frontal | R | 14 | 6 | 52 |
| Posterior-Medial Frontal | R | 8 | 14 | 60 |
| Middle Frontal Gyrus | L | -26 | 44 | 24 |
| Middle Frontal Gyrus | L | -30 | 34 | 34 |
| Middle Frontal Gyrus | L | -36 | 36 | 22 |
| Precentral Gyrus | R | 46 | 0 | 48 |
| Middle Frontal Gyrus | R | 36 | -4 | 52 |
| Middle Frontal Gyrus | R | 40 | 20 | 34 |
| Anterior Cingulate Cortex | L | -8 | 30 | 20 |
| Mid-Cingulate Cortex | L | -6 | 10 | 34 |
| Anterior Cingulate Cortex | L | -2 | 16 | 22 |
| Cerebellar Vermis | | 0 | -58 | -34 |
| Cerebellum | L | -10 | -44 | -14 |
| Superior Temporal Gyrus | L | -58 | -6 | 6 |
| Rolandic Operculum (Insula) | L | -46 | 4 | 8 |
| Heschls Gyrus | L | -52 | -16 | 8 |
| Precuneus | L | -8 | -56 | 18 |

| | | | | |
|---|---|---|---|---|
| Posterior Cingulate Cortex | L | -6 | -48 | 20 |
| Cerebellum | R | 28 | -52 | -22 |
| Cerebellum | R | 18 | -36 | -16 |
| Cerebellum | R | 18 | -44 | -24 |
| Inferior Frontal Gyrus (p. Triangularis) | R | 54 | 34 | 10 |
| Inferior Frontal Gyrus (p. Triangularis) | R | 52 | 34 | 0 |
| Inferior Frontal Gyrus (p. Triangularis) | R | 44 | 36 | 6 |

Table S5a: SSRT correlates for increasing SSRT for pNUD group at cue epoch

| Brain Region | Side | X | Y | Z |
|---|---|---|---|---|
| Anterior Cingulate Cortex | L | 0 | 36 | -8 |

Table S5b: SSRT correlates for increasing SSRT for pNUD group at anticipation epoch

| Brain Region | Side | X | Y | Z |
|---|---|---|---|---|
| Posterior-Medial Frontal | R | 12 | 6 | 52 |
| Mid-Cingulate Gyrus | L | -8 | 18 | 32 |
| Mid-Cingulate Gyrus | R | 6 | 16 | 42 |
| Inferior Frontal Gyrus (p. Triangularis) | L | -36 | 32 | 20 |
| Precentral | L | -52 | 2 | 28 |
| Insula Lobe | L | -30 | 14 | 6 |
| Superior Parietal Lobule | L | -22 | -58 | 52 |
| Inferior Parietal Lobule | L | -46 | -36 | 52 |
| Precentral Gyrus | R | 50 | 6 | 32 |
| Inferior Frontal Gyrus (p. Opercularis) | R | 44 | 4 | 22 |
| Precentral Gyrus | R | 60 | 4 | 34 |
| Precentral Gyrus | R | 38 | -2 | 42 |
| Precentral Gyrus | R | 46 | 0 | 48 |
| Superior Parietal Lobule | R | 18 | -56 | 54 |
| Inferior Parietal Lobule | R | 30 | -54 | 52 |
| Insula Lobe | R | 34 | 12 | 4 |
| Inferior Frontal Gyrus (p. Triangularis) | R | 44 | 20 | 6 |
| Insula Lobe | R | 30 | 26 | 4 |

Table S5c: SSRT correlates for increasing SSRT for pNUD group at trial epoch

| Brain Region | Side | X | Y | Z |
|---|---|---|---|---|
| Inferior Frontal Gyrus (p. Opercularis) | R | 54 | 20 | 34 |
| Inferior Frontal Gyrus (p. Triangularis) | R | 56 | 28 | 20 |
| Precentral Gyrus | R | 34 | 4 | 30 |
| Inferior Frontal Gyrus (p. Triangularis) | L | -56 | 22 | 16 |

| | | | | |
|---|---|---|---|---|
| Inferior Frontal Gyrus (p. Triangularis) | L | -56 | 14 | 30 |
| Inferior Frontal Gyrus (p. Opercularis) | L | -48 | 14 | 20 |
| Superior Medial Gyrus | L | -4 | 36 | 42 |
| Posterior-Medial Frontal | R | 8 | 18 | 50 |
| Superior Medial Gyrus | R | 10 | 38 | 52 |
| SupraMarginal Gyrus | L | -62 | -34 | 34 |
| SupraMarginal Gyrus | L | -54 | -42 | 30 |
| Inferior Parietal Lobule | L | -56 | -40 | 44 |
| Inferior Parietal Lobule | R | 54 | -36 | 46 |
| SupraMarginal Gyrus | R | 64 | -32 | 36 |
| Middle Frontal Gyrus | R | 48 | 46 | 16 |
| Inferior Frontal Gyrus (p. Triangularis) | R | 50 | 32 | 6 |

Table S5d: SSRT correlates for increasing SSRT for pNUD group at feedback epoch

| Brain Region | Side | X | Y | Z |
|---|---|---|---|---|
| Middle Occipital Gyrus | R | 34 | -90 | 10 |
| Middle Occipital Gyrus | R | 42 | -82 | 12 |
| Inferior Occipital Gyrus | R | 38 | -88 | -4 |
| Superior Parietal Lobule | R | 30 | -58 | 60 |
| Superior Parietal Lobule | R | 40 | -48 | 58 |
| Superior Parietal Lobule | R | 20 | -64 | 60 |
| Inferior Frontal Gyrus (p. Opercularis) | R | 54 | 18 | 34 |
| Inferior Frontal Gyrus (p. Triangularis) | R | 54 | 26 | 20 |
| Precentral Gyrus | R | 60 | 4 | 32 |
| Calcarine Gyrus | R | 24 | -66 | 6 |
| Calcarine Gyrus | R | 22 | -56 | 4 |
| Calcarine Gyrus | R | 16 | -62 | 8 |
| Inferior Frontal Gyrus (p. Opercularis) | L | -46 | 14 | 22 |
| Inferior Frontal Gyrus (p. Triangularis) | L | -56 | 14 | 30 |
| Middle Frontal Gyrus | L | -50 | 14 | 40 |
| Inferior Occipital Gyrus | L | -38 | -78 | -12 |
| Middle Occipital Gyrus | L | -34 | -86 | 2 |
| Middle Occipital Gyrus | L | -30 | -90 | 14 |
| Inferior Parietal Lobule | L | -32 | -58 | 56 |
| Superior Parietal Lobule | L | -24 | -62 | 48 |
| Superior Parietal Lobule | L | -22 | -66 | 40 |
| Posterior-Medial Frontal | R | 6 | -2 | 48 |
| Mid-Cingulate Cortex | L | -4 | -2 | 46 |
| Posterior-Medial Frontal | R | 14 | -6 | 54 |

Table S5e: SSRT correlates for decreasing SSRT for pNUD group at feedback epoch

| Brain Region | Side | X | Y | Z |
|---|---|---|---|---|
| Superior Occipital Gyrus | L | -14 | -90 | 2 |
| Superior Occipital Gyrus | L | -12 | -90 | 12 |
| Superior Occipital Gyrus | L | -20 | -94 | 20 |
| Precuneus | L | -12 | -42 | 4 |
| Mid Orbital Gyrus | R | 6 | 54 | -2 |
| Anterior Cingulate Cortex | L | -6 | 50 | -2 |
| Mid Orbital Gyrus | R | 10 | 42 | -2 |

Table S5f: SSRT correlates for increasing SSRT for control group at feedback epoch

| Brain Region | Side | X | Y | Z |
|---|---|---|---|---|
| Calcarine Gyrus | L | -6 | -90 | -4 |
| Calcarine Gyrus | R | 16 | -78 | 14 |
| Lingual Gyrus | R | 6 | -72 | 4 |
| Posterior-Medial Frontal | R | 8 | 10 | 48 |
| Posterior-Medial Frontal | R | 12 | 0 | 60 |
| Cerebellum | R | 12 | -66 | -16 |
| Cerebellar Vermis | R | 4 | -70 | -16 |
| Cerebellum | R | 16 | -72 | -20 |

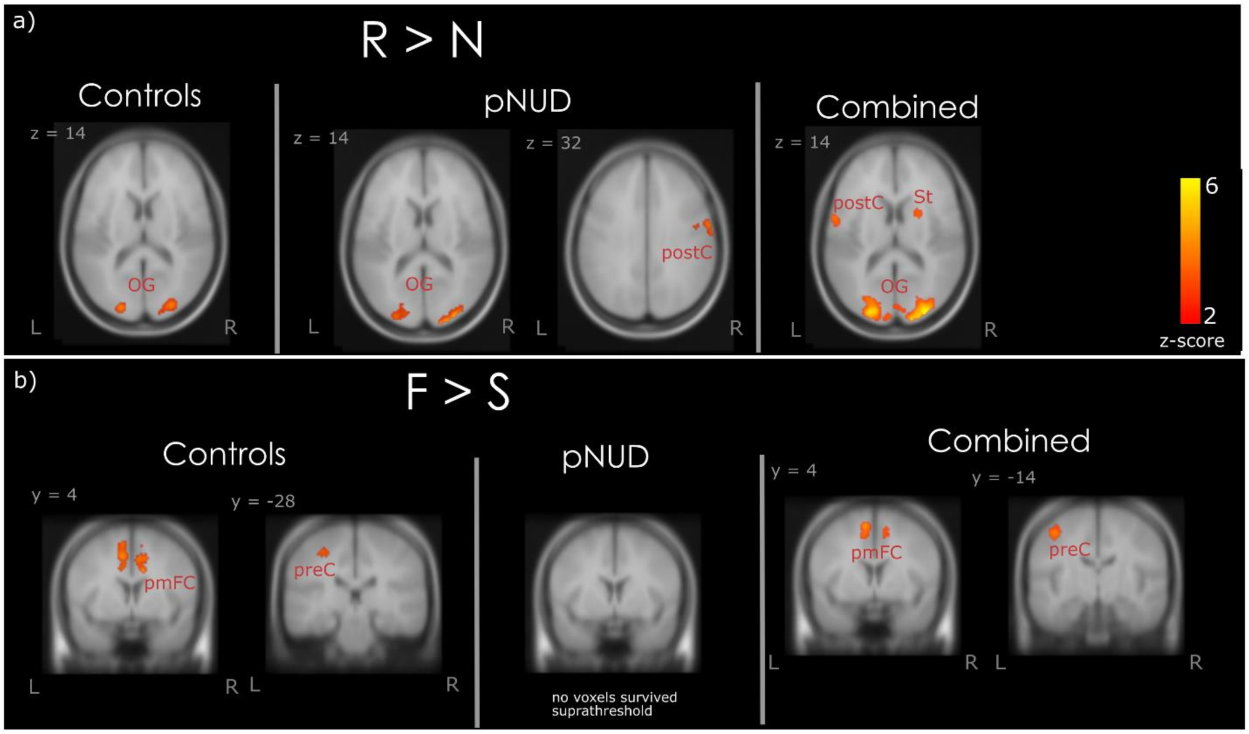


Figure S2. *Activity during cue epoch.* This is activity when participants are cued on the probability of an upcoming stop trial and whether it is a reward or a neutral trial. *Abbreviations: R = reward, N = neutral, F = Failed inhibition, S = successful inhibition, pNUD = people with a nicotine disorder group, C = control group. PC = precuneus, St = striatum, pmFC = posterior medial frontal corte, preC = precentral, postC = postcentral, OG = occipital gyrus. L = left, R = Right.* *subthreshold left caudate (MNI: -40, -34, -2) activated for R>N contrast for controls at p = 0.088 (not shown in figure as it is subthreshold).

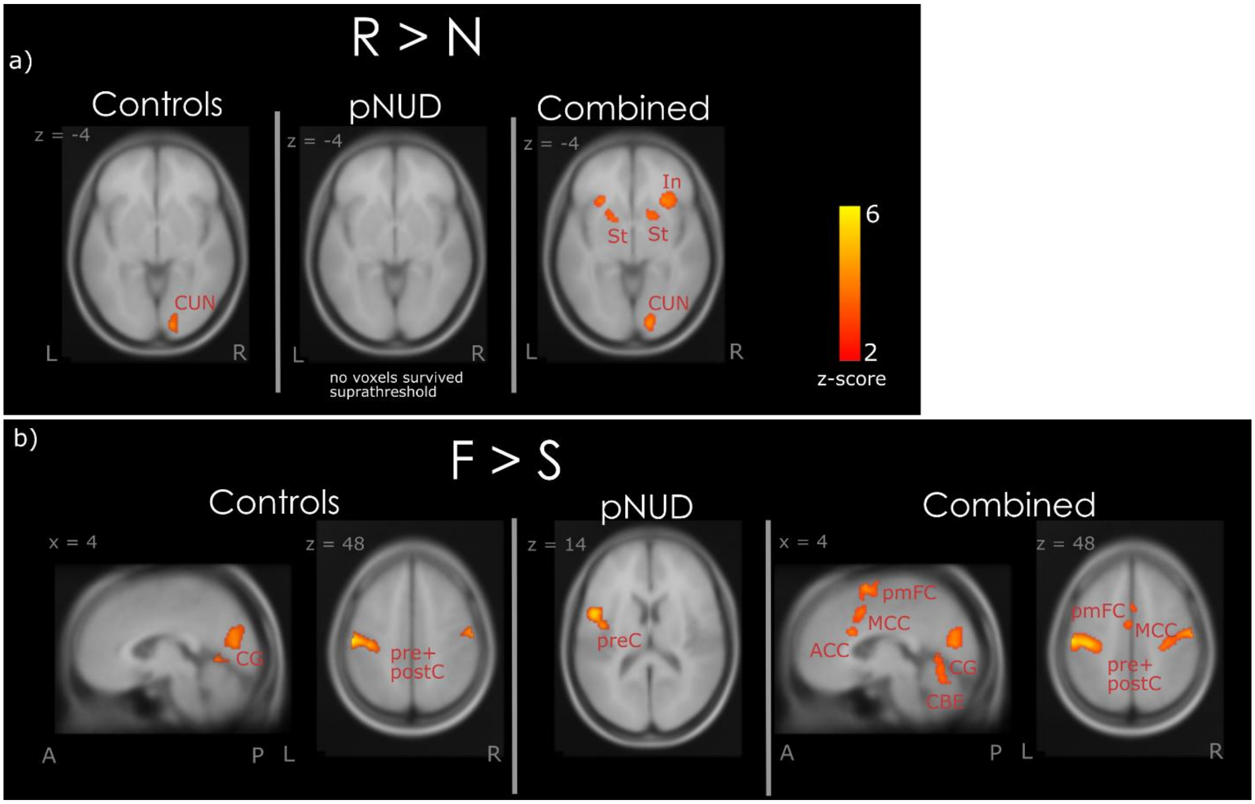


*Figure S3. Brain activity during the anticipation epoch.* This is activity immediately prior to the onset of the trial. *Abbreviations: R = reward, N = neutral, F = Failed inhibition, S = successful inhibition, pNUD = people with a nicotine disorder group, C = control group, RS = reward successful inhibition trials, NF = neutral failed inhibition trials In = insula, St = striatum, CUN = cuneus, ACC = anterior cingulate cortex, MCC = midcingulate cortex, pmFC = posterior medial frontal cortex, preC = precentral, postC = postcentral, IFG = inferior frontal cortex, SMG = supramarginal gyrus, MFG = middle frontal cortex, IPL = inferior parietal lobule, CBE = cerebellum. L = left, R = Right. A = anterior, P = posterior.*

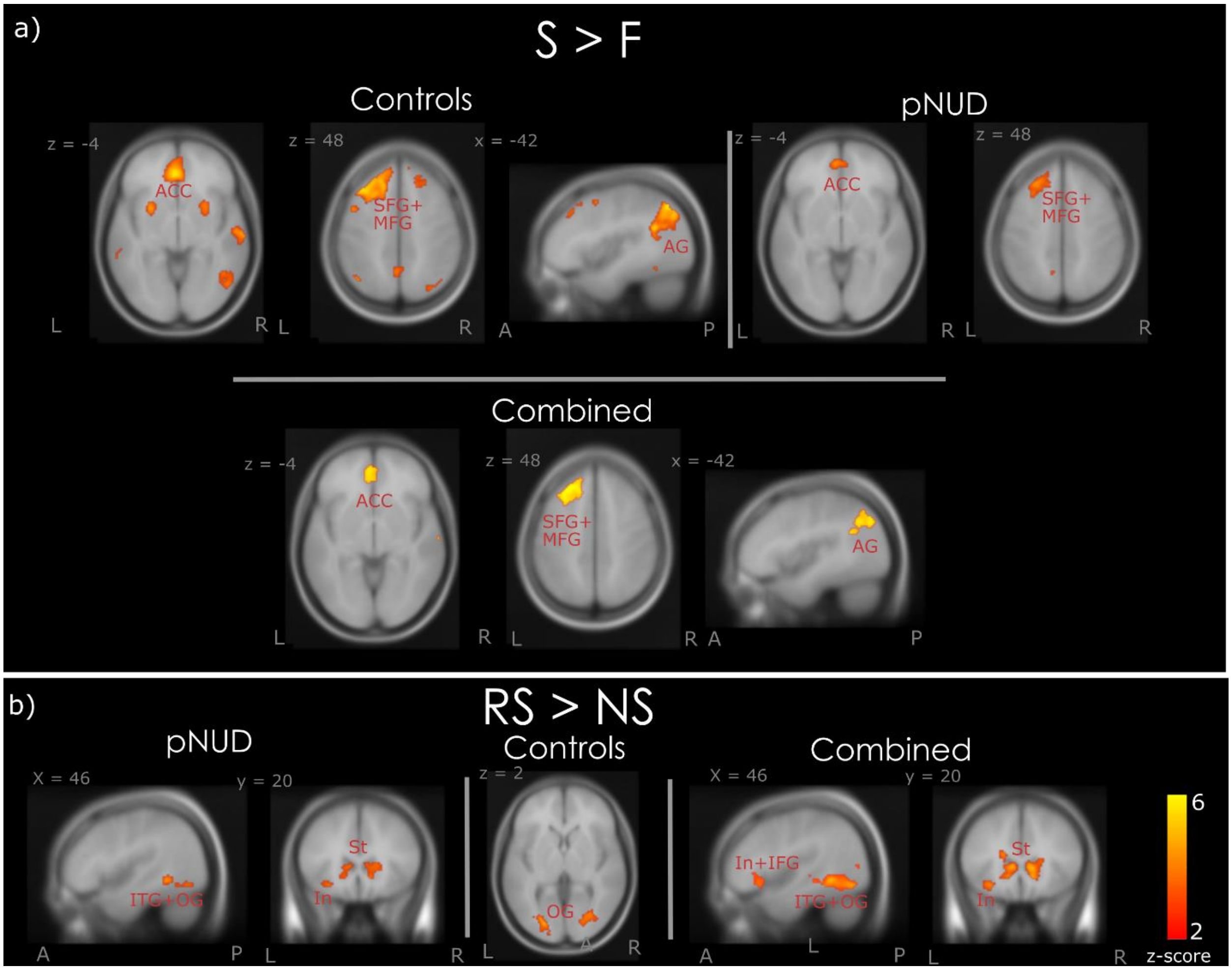


Figure S4. *Brain activity at the stop-trial epoch.* This is activity at the trial – when the stop signal is presented, and participants need to inhibit a response to make progress towards the larger later reward. *Abbreviations: R = reward, N = neutral, F = Failed inhibition, S = successful inhibition, RS = reward successful inhibition trials, NS = neutral successful inhibition trials, pNUD = people with a nicotine disorder group, C = control group. ACC = anterior cingulate cortex, SFG = superior frontal gyrus, MFG = middle frontal gyrus, In = insula, IFG = inferior frontal gyrus, St = striatum, AG = angular gyrus, PL = parietal lobule. L = left, R = Right. A = anterior, P = posterior.*

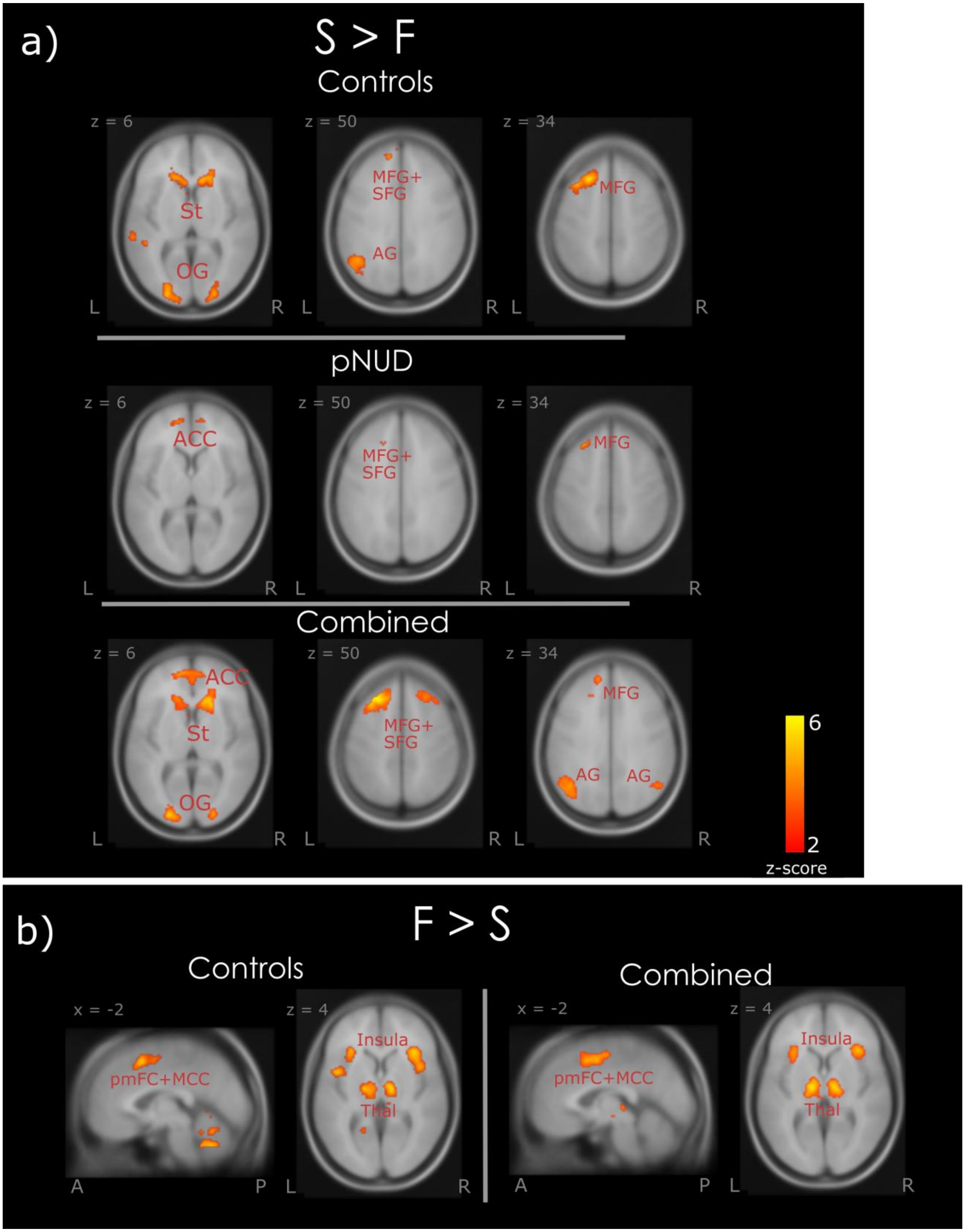
a)
S > F
Controls
z = 6
St
OG
z = 50
MFG+
SFG
AG
z = 34
MFG
pNUD
ACC
MFG+
SFG
MFG
Combined
ACC
St
OG
MFG+
SFG
MFG
AG
AG
6
2
z-score
b)
F > S
Controls
x = -2
pmFC+MCC
z = 4
Insula
Thal
Combined
x = -2
pmFC+MCC
z = 4
Insula
Thal
L
R
A
P

Figure S5. *Brain activity at the feedback epoch.* This is activity at the feedback epoch, where participants are given feedback of “correct 0c” for successful inhibitions and “incorrect 0c” for failed inhibitions, where the response was slower than 400ms or was the incorrect button press. *Abbreviations: R = reward, N = neutral, F = Failed inhibition, S = successful inhibition, RS = reward successful inhibition trials, NS = neutral successful inhibition trials, pNUD = people with a nicotine disorder group, C = control group. ACC = anterior cingulate cortex, SFG = superior frontal gyrus, MFG = middle frontal gyrus, In = insula, IFG = inferior frontal gyrus, MCC = midcingulate gyrus, PCC = posterior cingulate cortex, pmFC = posterior medial frontal cortex, OG = occipital gyrus, St = striatum, Thal = thalamus, CBE = cerebellum, SMG = superior medial gyrus, AG = angular gyrus. L = left, R = Right. A = anterior, P = posterior.*
